# PII interactions with BADC and BCCP proteins co-regulate lipid and nitrogen metabolism in Arabidopsis

**DOI:** 10.1101/2024.11.04.621944

**Authors:** Matthew G. Garneau, Gabriel Lemes Jorge, Jay Shockey, Jay J. Thelen, Philip D. Bates

**Affiliations:** Institute of Biological Chemistry, Washington State University, Pullman, Washington 99164, United States; Department of Biochemistry and Christopher S. Bond Life Sciences Center, University of Missouri, Columbia MO, 65221, United States; United States Department of Agriculture, Agricultural Research Service, Southern Regional Research Center, New Orleans, Louisiana 70124, United States

## Abstract

In plants the initiation of fatty acid synthesis is catalyzed by acetyl-CoA carboxylase (ACCase) which produces malonyl-CoA. The heteromeric form of ACCase (htACCase) is a holoenzyme consisting of biotin carboxylase and carboxyltransferase sub-complexes, both of which are subject to extensive regulation. Biotin carboxylase activity is controlled in part by the presence of the catalytic biotin carboxyl carrier proteins (BCCP1/2) and/or the non-catalytic, non-biotinylated, biotin/lipoyl attachment domain-containing proteins (BADC1/2/3) that associate with backbone biotin carboxylase (BC) protein. However, the mechanisms regulating BADC and BCCP interaction with BC and thus ACCase activity *in planta* are not clear. Here we demonstrate the *Arabidopsis thaliana* regulatory protein PII modulates htACCase activity through independent interactions with BADC and BCCP proteins in a selective manner. Analysis of *badc1/2* and *badc1/3* mutant lines and the respective *pii* triple mutants reveal that changes in seed oil and protein accumulation of *badc* double mutants are PII/nitrogen dependent. Absolute quantification of htACCase subunits and PII in developing seeds suggests that Arabidopsis exerts tight regulation over individual protein stoichiometry to balance oil and protein accumulation. The effects on vegetative and seed development indicate PII and BADC proteins have distinct but overlapping roles in the regulation of plant metabolism.

## Introduction

Seeds are amongst the highest value products produced by commercial agriculture, and breeding efforts have focused on improving both seed yield and quality. Seed crops from the Brassicaceae family in particular are valued for their oil and protein accumulation, which can be used for food, biofuel, and animal feed (Raboanatahiry et al., 2021; So and Duncan, 2021). However, efforts to genetically improve seed oil content have largely indicated that the ratio of seed oil to protein is generally inversely related (Vigeolas et al., 2007; Schwender and Hay, 2012; Jasinski et al., 2018; Wang et al., 2020; Wang et al., 2022). One possible reason for the inverse relationship between oil and protein is that the metabolic precursors for amino acids and fatty acids that make up protein and oil, respectively, are in competition for the same pool of carbon supplied by the parent plant (Schwender et al., 2003; Schwender and Hay, 2012; Rolletschek et al., 2020; Song et al., 2023). Therefore, coordinated metabolic regulation of oil and protein biosynthesis has likely evolved to establish their relative contribution. Seeds have developed a complex regulatory framework in response to the competition between carbon precursors that takes into account nitrogen (N) status, biotic and abiotic stress, cell determination, and a host of other factors (Su et al., 2021; Yang et al., 2022). Yet many aspects of the regulatory cross-talk between oil and protein metabolism in seeds are not yet fully understood.

There is ample evidence that enhanced seed oil production can be achieved by increasing fatty acid synthesis, fatty acid transport, and/or lipid assembly (Baud, 2018; Sagun et al., 2023). *In vivo* modulation of total fatty acid and oil synthesis has been largely attributed to the regulation of acetyl-CoA carboxylase (ACCase) which catalyzes the production of malonyl-CoA from acetyl-CoA and carbonate, and is the first committed step of fatty acid biosynthesis (Li-Beisson et al., 2013). Plants contain homomeric and heteromeric forms of ACCase, but it is the heteromeric form (htACCase) that confers modular protein complex assembly, and for which multifaceted regulation is known. The htACCase is regulated by a large number of factors including redox and pH status of the stroma, protein phosphorylation, feedback inhibition, substrate availability, and most recently two gene families of effector proteins (Huerlimann and Heimann, 2013; Salie and Thelen, 2016; Ye et al., 2020b; Ye et al., 2020a). Eukaryotic algae and dicotyledonous plants initiate fatty acid synthesis in the plastid through the activity of htACCase which includes both catalytic and non-catalytic regulatory proteins. Plant htACCase is composed of two catalytic sub-complexes, biotin carboxylase and carboxyltransferase complexes (Salie and Thelen, 2016). In the Brassicaceae oilseed *Arabidopsis thaliana* the functional biotin carboxylase sub-complex is comprised of biotin carboxylase (BC), and at least one of two biotin carboxyl carrier proteins (BCCP1/2) or a BCCP and one of three biotin/lipoyl attachment domain containing protein (BADC1/2/3) which are ancient orthologs of BCCP but are not biotinylated (Salie and Thelen, 2016). The biotin carboxylase complex catalyzes ATP-dependent carboxylation of the biotin moiety which is then utilized by the carboxyltransferase complex to carboxylate acetyl-CoA to malonyl-CoA. The carboxyl transferase complex consists of a heterotetramer of α and β-carboxyltransferases (α/β-CT). Plant htACCase has been hypothesized to have a similar subunit stoichiometry to that of *E. coli* which consists of two biotin carboxylase complexes for each carboxyltransferase complex [(BCCP/BADC)_4_(BC)_2_(α-CT)_2_(β-CT)_2_], however, this has yet to be experimentally verified in plants (Salie and Thelen, 2016; Cronan, 2021). The modulation of the complexity of htACCase holoenzyme assembly appears to be an important part of the regulation of *de novo* fatty acid synthesis.

Beyond htACCase regulation through expression, redox, pH and substrate availability; recent results suggest that assembly of the holoenzyme (including interacting regulatory proteins) is key to fine tuning htACCase activity. The carboxyltransferase complex is regulated by the stoichiometric levels of α-CT (Wang et al., 2022) as well as its anchoring to the plastid membrane by CTI proteins (Ye et al., 2020a). In addition, there is mounting evidence that ACCase activity is in part controlled through the relative availability of BADC and BCCP subunits for biotin carboxylase complex assembly. BADCs are homologous to BCCP proteins and both contain folded structured domains and large intrinsically disordered regions, however, BADCs do not contain the conserved lysine required for attachment of the biotin cofactor required for catalytic activity (Ye et al., 2020b). BADC proteins were initially described and subsequently characterized as negative regulators (Salie et al., 2016b; Keereetaweep et al., 2018; Ye et al., 2020b), however, recent evidence also suggests that the roles of BADC proteins may be more dynamic. *In vitro* measurements of BADC and BCCP interaction with the backbone BC protein demonstrated that pH, cationic conditions, and phosphate availability all affected protein binding affinity (Ye et al., 2020b). Reconstituted protein assays of htACCase have also hypothesized that BADCs are involved in the formation of the active holoenzyme quaternary structure (Shivaiah et al., 2020). Additionally, BADCs are transcriptionally regulated by the WRI1 transcription factor, a master regulator of oil accumulation, and BADC expression in plant seeds is upregulated during developmental stages of rapid fatty acid biosynthesis and oil accumulation (Liu et al., 2019). There is also evidence of circadian regulation of BADC2 and BCCP2, however, the direct regulatory pathway has yet to be understood (Kim et al., 2023).

Regardless of the exact function of individual BADCs as negative regulators and/or holoenzyme assembly factors within the ACCase complex, physiological studies have shown that *badc1/2* and *badc1/3* double mutants have large effects on plant development. Arabidopsis *badc1/2* knockout lines have reduced root and shoot development, possibly through changes in auxin catabolism (Keereetaweep et al., 2018; Liu et al., 2019). Seeds of both *badc1/2* and *badc1/3* mutations exhibit increased seed oil content while the *badc2/3* combination is embryo-lethal (Keereetaweep et al., 2018; Shivaiah et al., 2020; Ye et al., 2020b). The *badc1/3* double mutant seed oil phenotype is also dependent on circadian regulation, likely as a result of light-induced changes to the stroma pH and ionic milieu leading to confirmation changes of BADC and BCCP proteins which alters their affinity to BC (Ye et al., 2020b), and demonstrates their ability for dynamic regulation based on the cellular metabolic status. Additional multi-omics analysis of developing *badc1/3* seeds have shown global development changes including altered fatty acid synthesis and turnover, as well as enhanced seed storage protein content (Kataya et al., 2024). Despite substantial prior research, the complex nature of BADC regulation including their interaction with biotin carboxylase, effects on fatty acid biosynthesis, and the coordination of fatty acid biosynthesis with protein synthesis are not yet fully understood.

In addition to the regulatory CTI and BADC proteins, there is evidence that ACCase is activity is co-regulated by the PII protein. PII is a highly conserved effector protein found in bacteria, algae, and plants. It is generally accepted that PII forms an active homo-trimer that can interact with and regulate C/N metabolism during abundant cellular energy states with a high ratio of ATP/ADP, and high N conditions that correspond to lower levels of 2-oxoglutarate (2-OG), which is consumed during N assimilation (Gerhardt et al., 2020a; Selim et al., 2020). In bacteria, PII has been demonstrated to measure both cellular energy status and N status through competitive binding of ATP (± Mg^2+^) or ADP, and 2-OG at metabolite binding sites which form at the lateral clefts between PII subunits (Fokina et al., 2010; Truan et al., 2010). Substrate binding to these sites introduce structural changes in PII’s interaction domain (called the T-loop) which regulates PII binding to other proteins (Smith et al., 2003; Fokina et al., 2010; Truan et al., 2010; Fokina et al., 2011; Gerhardt et al., 2020b; Selim et al., 2020). Bacterial PII is also regulated through phosphorylation which has not yet been found in higher plants and may represent a major difference between plant and bacterial PII regulation (Smith et al., 2004). In plants, active PII directly regulates the activity of N-acetyl glutamate kinase (NAGK), which catalyzes the second committed step in ornithine and arginine biosynthesis (Feria Bourrellier et al., 2009). Arabidopsis PII activity has further been described as regulating nitrite import into the chloroplast (Ferrario-Méry et al., 2005; Ferrario-Méry et al., 2008). However, the direct protein targets of PII interaction have yet to be identified (Hsieh et al., 1998; Ferrario-Méry et al., 2005; Ferrario-Méry et al., 2008). Interestingly, PII’s role in N metabolism is glutamine-dependent in all higher plants except for *Brassicaceae* (Smith et al., 2003; Ferrario-Méry et al., 2006; Feria Bourrellier et al., 2009). PII regulation of fatty acid synthesis is also highly conserved from bacteria to higher plants. In Arabidopsis, prior research suggests that active PII (high energy/ N status) binds to BCCP proteins as a negative regulator of fatty acid synthesis (Feria Bourrellier et al., 2010b). However, in *pii* mutants, the loss of PII only conditionally affects seed oil and protein composition, suggesting a role in fine-tuning plastid fatty acid synthesis to correlate with fatty acid elongation in the cytosol (Baud et al., 2010). There is also limited evidence that PII may directly interact with BADC proteins (Baud et al., 2009).

To ascertain whether the combined loss of negative regulators of biotin carboxylase (BADCs and PII) could be a strategy to improve seed oil content, *pii/badc1/2* and *pii/badc1/3* triple mutants were produced and analyzed. Contrary to our initial hypothesis, the results demonstrate that the enhanced ACCase activity and seed oil content of *badc* double mutants is dependent on the presence of PII, and that PII plays a role in modulating the activity of ACCase in response to energy and N status through selective interactions with both BADC and BCCP subunits. Together the results presented herein indicate more complex roles for PII and BADC than simple negative regulators, and suggest they work together to regulate ACCase activity in response to cellular energy and N status.

## Results

### Differential effects of *pii* and *badc* mutant combinations on vegetative tissue development

Both traditional crossing and CRISPR/Cas9 based genome editing approaches were used to generate triple loss-of-function mutants lacking functional *PII* and *BADC1* and either *BADC2* or *BADC3*. The first successful lines produced were the T-DNA cross for the *pii/badc1/2* triple mutant generated by the cross of the *pii* and *badc1/2* tDNA insertion lines, and simultaneous CRISPR/Cas9 mutation of *badc1* and *badc3* in the *pii* T-DNA background. These lines were used for all subsequent experiments. Successively, the *pii/badc1/3* T-DNA cross was obtained and had the same oil phenotype as the line containing CRISPR/Cas9 based alleles (Fig. S1). Since PII and BADC proteins were previously described as negative regulators of ACCase, we hypothesized the triple mutants could have unregulated ACCase, resulting in increased fatty acid biosynthesis. Over production of free fatty acids (those not incorporated into glycerolipids) is cytotoxic, thus we first analyzed the effect of each mutant combination on vegetative growth. None of the mutant lines displayed necrotic lesions associated with free fatty acid induced cytotoxicity (Fan et al., 2013), suggesting no overproduction of free fatty acids (Fig. 1A). The *badc1/2* double mutant was significantly smaller than WT, while *badc1/3* and *pii* mutants showed no significant change (Fig 1A, B), consistent with prior results (Ferrario-Méry et al., 2005; Keereetaweep et al., 2018). Loss of *pii* in both *badc1/2* and *badc1/3* backgrounds led to a small but significant decrease in overall rosette size (Fig. 1A) and rosette biomass (Fig. 1B). Root growth in 10-day old Arabidopsis seedlings was also analyzed. As reported previously, the *badc1/2* seedling roots were significantly smaller than WT (Fig. 1C, D). Interestingly, while both the *pii* mutant and *badc1/3* double mutant roots were significantly increased in length, the loss of *pii* in each of the *badc* backgrounds led to a trend of decreased average root length that was not significantly different than either *badc1/3* and WT (Fig. 3D). Therefore, stacking the *pii* mutation into the *badc1/2* or *badc1/3* backgrounds leads to a consistent small reduction in shoot and root biomass.

**Figure 1:**
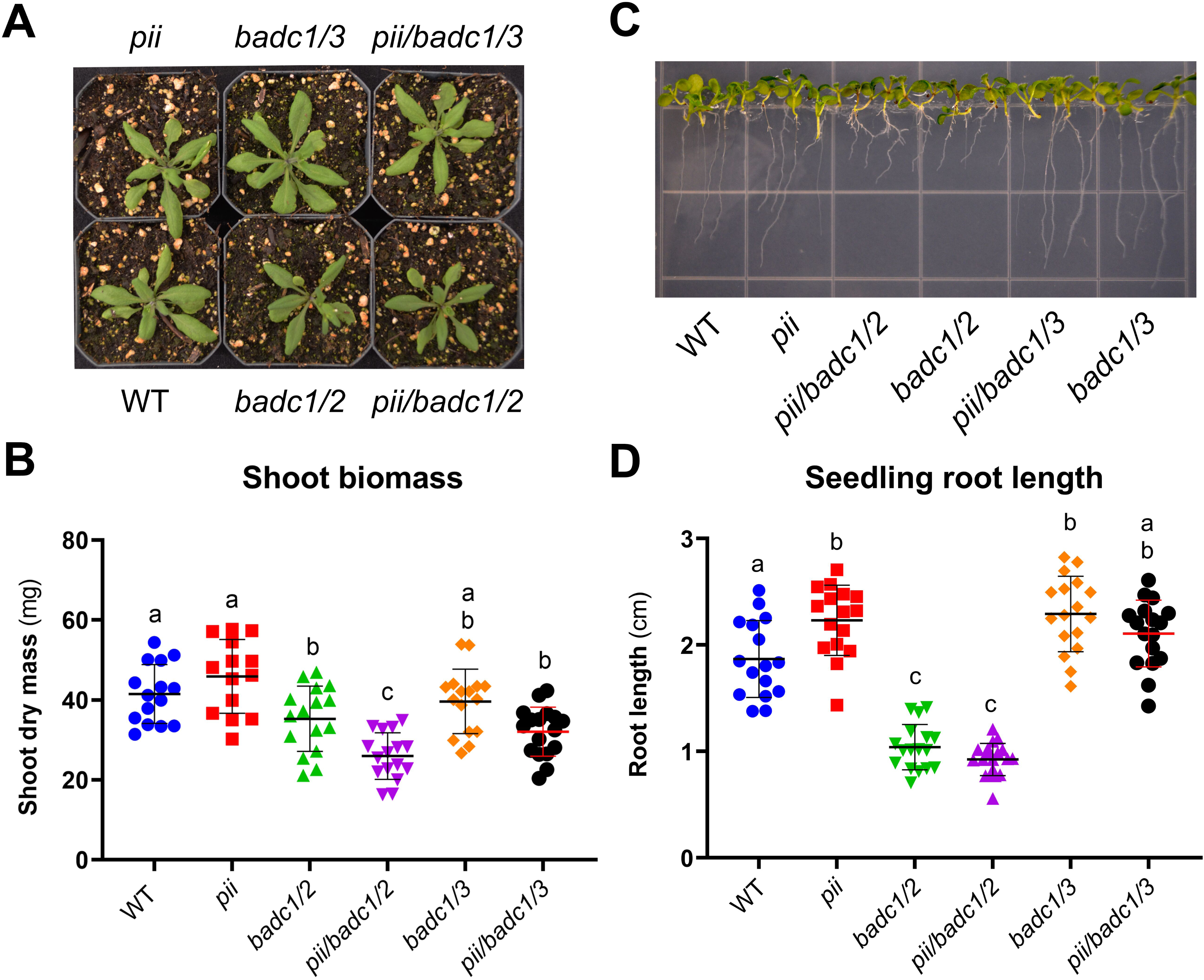
Vegetative tissue biomass of WT and mutant lines. Arabidopsis wild type (WT), mutants of *pii, badc1/2, badc1/3,* and their crosses were analyzed for shoot area (A) and biomass (B) in 3-week-old plants. Root development was analyzed in 10-day-old seedlings (C) and average root lengths were compared (D). Data is presented as mean ± standard deviation (SD), statistical differences were analyzed by ANOVA and Holm-Šidák multiple comparison test, different letters above each line represent significant differences in means (p-value < 0.05).

### Stacking *pii* into both *badc* double mutants reduces seedling ACCase activity, but the effect on leaf lipid and protein content is dependent on the *badc* mutant background

To investigate the effect of *badc* and *pii* mutations on leaf metabolism, the rosette leaves of well-fertilized Arabidopsis were analyzed for changes in lipid and protein content. Only the lipid content in *badc1/2* was significantly different from WT with a ∼7% decrease; interestingly WT lipid content was recovered in the *pii/badc1/2* triple mutant (Fig. 2A). To determine if changes in leaf lipids were altered by potential changes in N metabolism, protein (the highest proportional nitrogenous component) was assayed. Leaf protein increased by 15% in the *badc1/2* double mutants (Fig. 2B). Similar to the leaf lipid content, protein levels in *pii/badc1/2* leaves returned to WT levels. Thus, changes in leaf *badc1/2* lipid content may be a consequence of increased leaf protein. In contrast, when htACCase activity was measured in developing seedlings, the *badc1/2* and *badc1/3* double mutants had modest but insignificant increases in average htACCase activity compared to WT. However, the absence of *pii* in either *badc1/2* and *badc1/3* backgrounds led to 46% and 38%, decreases in htACCase activity, respectively. When ACCase activity was measured per µg leaf protein similar trends were seen as for ACCase activity per seedling (Fig. S2). This result is surprising in that loss of the potential negative regulator PII decreased rather than increased seedling ACCase activity. In leaves of the *badc1/2* double mutant the decreased lipid accumulation may be the result of altered N metabolism through prioritization of carbon for protein synthesis, upregulation of fatty acid turnover, or both.

**Figure 2:**
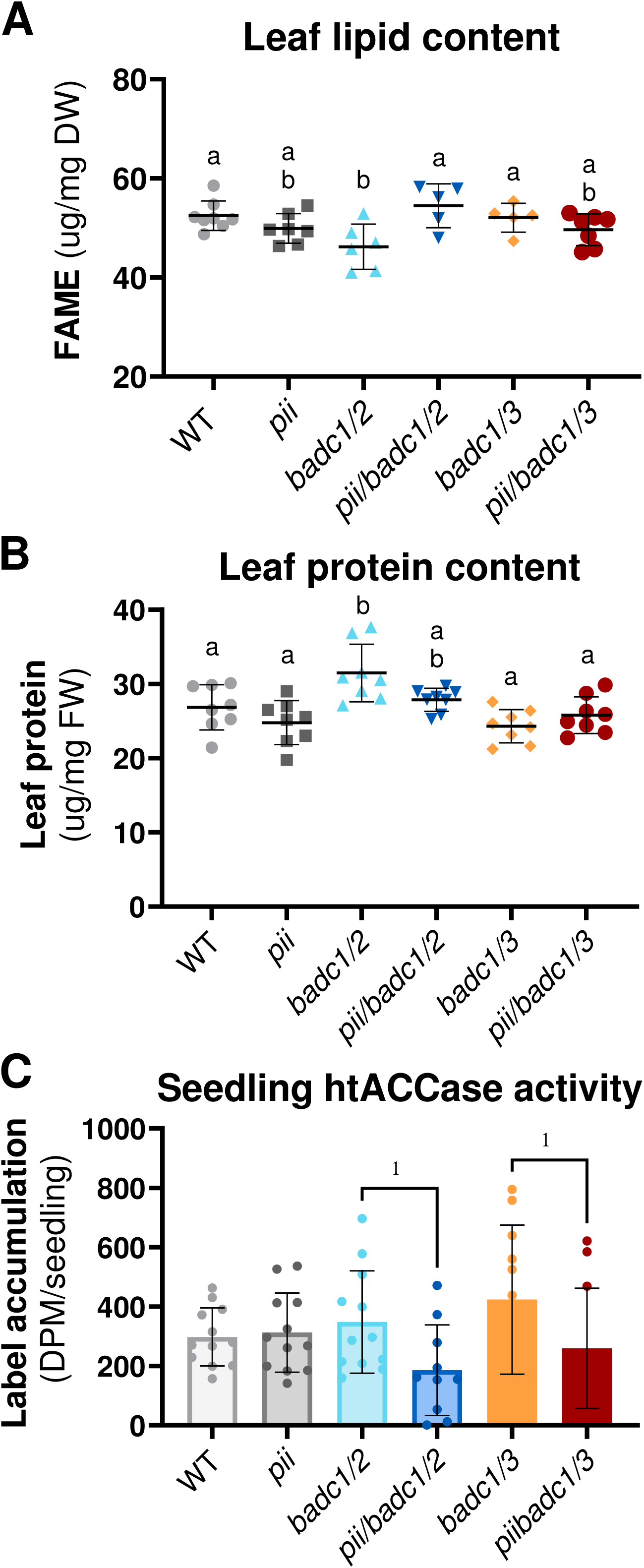
Leaf lipid, protein, and htACCase activity content of WT and mutant lines. Arabidopsis wild type (WT) and mutants of *pii, badc1/2, badc1/3,* and the triple mutant lines were analyzed for leaf oil content (A) and protein (B) in 3-week-old plants. The activity of HtACCase was measured in 10-day-old seedlings (C) by measuring the incorporation ^14^C-bicarbonate. Data is presented as mean ± SD, statistical differences were analyzed by ANOVA and Holm-Šidák multiple comparison tests for leaf oil and protein content. Different letters above each line represent significant differences in means (p-value < 0.05). Significant differences in ACCase activity between *badc* double and *pii/badc* triple mutants were analyzed by Welches t-test(*, *p*< 0.05).

### Leaf lipid composition is altered in *badc* double mutants and dependent on *pii* function

To further determine what effect each mutant combination may have on leaf lipid metabolism, lipids were extracted, separated by HPLC, derivatized to fatty acid methyl esters (FAMEs), and were quantified by GC-FID. Initial analysis of total fatty acid content of extracted leaf lipids indicated that both *badc* double mutants had increased levels of saturates (16:0, 18:0) and 7,10,13-hexadecatrienoic acid (16:3) at the cost of α-linolenic acid (18:3) as compared to WT (Fig. 3A). The stacking of *pii* into the *badc1/2* background resulted in the largest decrease in 18:3 with corresponding increases in oleic acid (18:1) and 16:0 (Fig. 3A). The decrease in fatty acid content in *badc1/2* leaves (Fig. 2A) was primarily due to reduced MGDG and PC content (Fig. 3B). Interestingly, the *pii/badc1/2* triple mutant largely returned membrane lipid contents back to WT (Fig. 3B). The fatty acid composition of MGDG and DGDG is in Fig. 3C and 3D, respectively, while the remaining lipids are in supplemental Figure S3. Analysis of MGDG and DGDG fatty acid composition demonstrated an increase in 16:3 and a corresponding decrease in 18:3 in the *badc* double mutant and *pii* triple mutant lines (Fig. 3C, 3D). Notably, although *badc1/2* exhibited a more pronounced leaf lipid phenotype, both *badc1/2* and *badc1/3* contained changes in thylakoid lipid ratios of 16:3/18:3, suggesting that both mutant backgrounds seem to have partially overlapping functions in vegetative tissue. Considering that 16:3 biosynthesis is through the chloroplast-localized prokaryotic pathway, the change in 16:3/18:3 ratio is consistent with relatively more galactolipids from the prokaryotic pathway rather than the eukaryotic pathway. This may suggest increased fatty acid biosynthesis in the plastid leading to higher acyl flux through the prokaryotic pathway and/or reduction of eukaryotic pathway galactolipid synthesis possibly due to turnover of excess fatty acids exported from the plastid. Therefore, the amount of leaf lipids and protein in *badc1/2* was dependent on the presence of PII, but the change in galactolipid fatty acid content was independent of PII presence and similar within both *badc* double mutant backgrounds.

**Figure 3:**
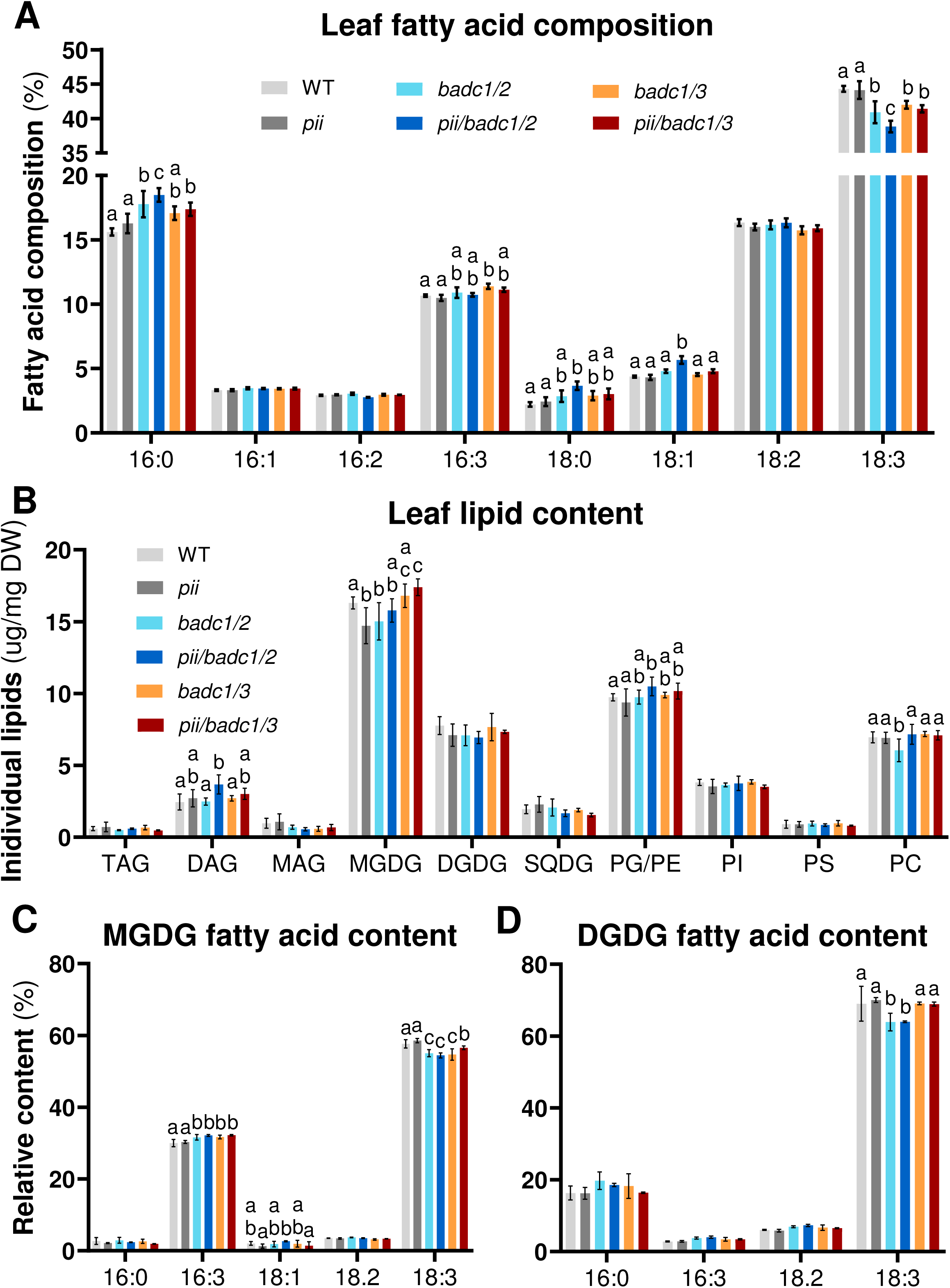
Analysis of leaf lipid and fatty acid composition in WT and mutant lines. Leaves of 3-week-old Col-0 wild type (WT), mutants of *pii, badc1/2, badc1/3,* and triple mutant lines were analyzed for changes in leaf total fatty acid composition (A) by GC-FID analysis of extracted leaf lipids (n=5). Individual leaf lipids were further separated by HPLC and resultant and composition of individual lipids was determined by GC-FID (B, n=3). The fatty acid content of MGDG (C) and DGDG (D) were analyzed and show changes in composition consistent with total leaf lipids (*n*=3). Data is presented as mean ± SD, statistical differences were analyzed by ANOVA and Holm-Šidák multiple comparison test, different letters above each line represent significant differences in means (p-value < 0.05).

### The *Badc1/2* line has increased nitrite sensitivity

Previous analysis of Arabidopsis *pii* mutants suggested an increase in the import of nitrite into the chloroplast that alters N metabolism in response to cellular energy status. While the mechanism of action has yet to be fully explained, possible protein:protein interactions between PII and BADC and/or BCCP can affect carbon demand from shared oil/protein precursor pools, may also contribute to the regulation of N metabolism. To investigate if the loss of BADCs also affects the ability of PII to regulate N metabolism, nitrite utilization was assayed in *pii, badc* double mutants, and their respective triple mutants through growth on media containing 10 mM nitrite as the sole N source (Fig. 4A). Nitrite is toxic unless reduced to ammonia in the chloroplast. The loss of functional PII is thought to enhance chloroplast nitrite import and detoxification (Oke, 1966; Ferrario-Méry et al., 2008; Krapp, 2015). Three-week-old seedlings that were green and had developed past the 2-cotyledon stage were counted as surviving (non-arrested seedlings) (Fig. 4B). The *pii* mutant demonstrated a 27% increase in seedling survival when only compared to WT, which is consistent with previously published results. Seedlings of *badc1/2* mutants had significantly less survivability with a 38% decrease compared to WT consistent with altered N transport/metabolism. Similar to *pii*, the corresponding triple mutant (*pii/badc1/2*) had increased survivability compared to the *badc1/2* background (Fig. 4B). In contrast, both *badc1/3* and *pii/badc1/3* were unaffected compared to *pii* or WT (Fig. 4B). Along with changes in leaf protein for *badc1/2* (Fig. 2B), these results suggest that the *badc1/2* mutant has a greater effect on N metabolism than *badc1/3*, possibly through the altered interaction of PII with the remaining BADC isoform in each mutant background.

**Figure 4:**
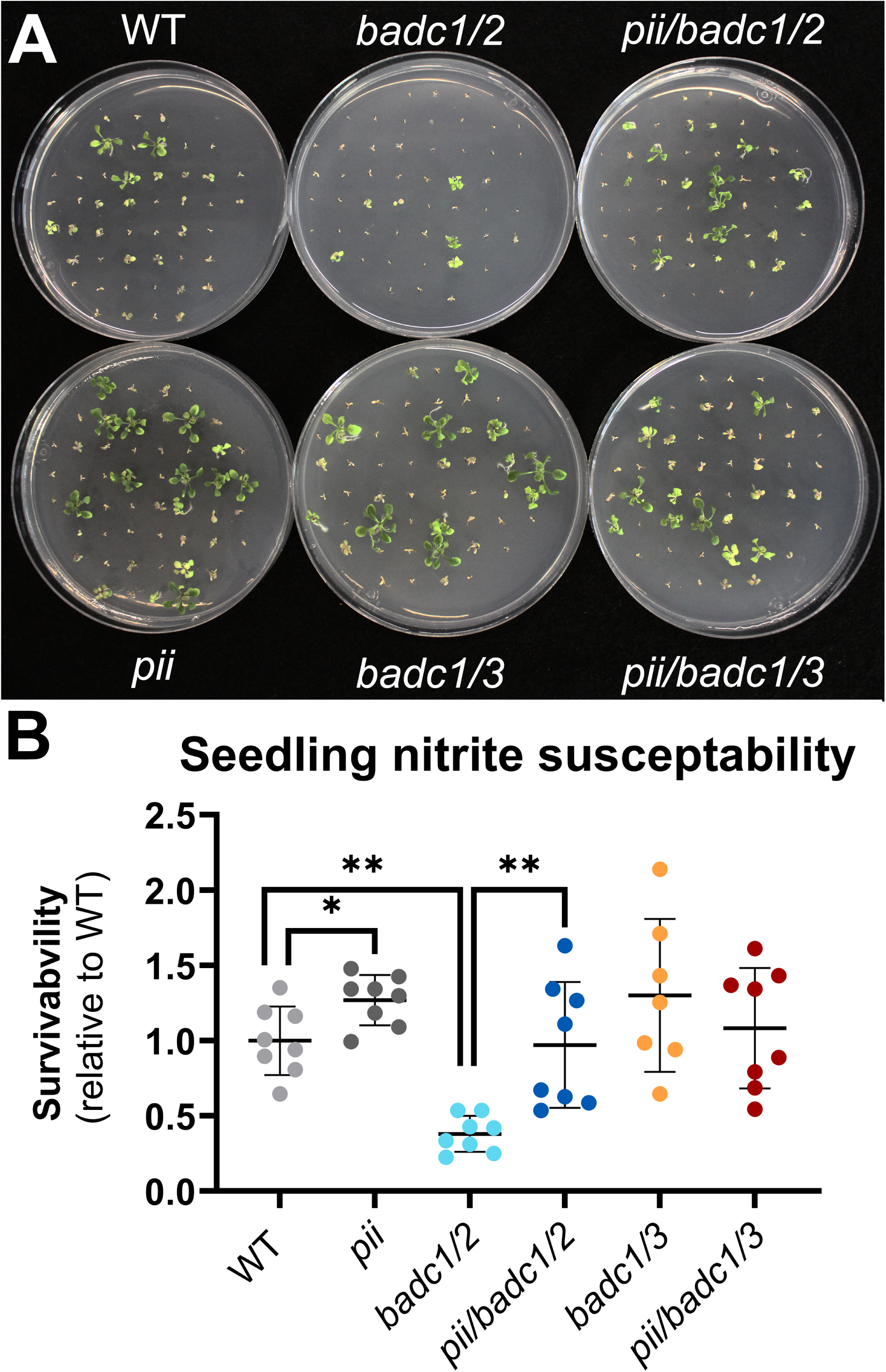
Determination of nitrite susceptibility in seedlings of badc1/2 and badc1/3 double mutants their respective pii/badc triple mutants. Arabidopsis Col-0 wild type (WT), mutants of *pii*, *badc1/2*, *badc1/3*, and triple mutant lines were geminated and grown on plates containing MS minimal media containing 10 mM nitrite as a sole nitrogen source. Plants were grown for 3 weeks and imaged (A). Seedling survival rate was calculated comparing the mean survival rate between WT and mutants on individual plates from 2 independent experiments (B, n=8). Data is presented as mean ± SD, significant differences to WT or ± *pii* were analyzed by Welches t-test (*, p< 0.05;**, p< 0.01).

### Seed oil and protein composition are altered in *badc1/2* and *badc1/3* lines in a PII and N fertilization dependent manner

To determine what effect the concurrent loss of *pii*, *badc1/2*, and *badc1/3*, and their respective triple mutants have on seed metabolism, both seed oil and protein were analyzed. Further, the role of PII in the regulation of *badc* double mutant lines in response to N was investigated by fertilizing plants using a modified Hoagland solution with either excess N (100 mg/plant) or no added nitrogenous compounds. Seeds of the well-fertilized (high N) *badc1/2* and *badc1/3* plants showed 12% and 6% increases in total fatty acid content, respectively, while mature *pii* seeds were not different than wild type (Fig. 5A), consistent with prior results (Baud et al., 2010; Keereetaweep et al., 2018). Interestingly, the loss of *pii* in either of the *badc* double knockout backgrounds reverted seed oil content back to WT levels (Fig. 5A), which supports that the *badc* seed oil phenotype is *pii* dependent.

**Figure 5:**
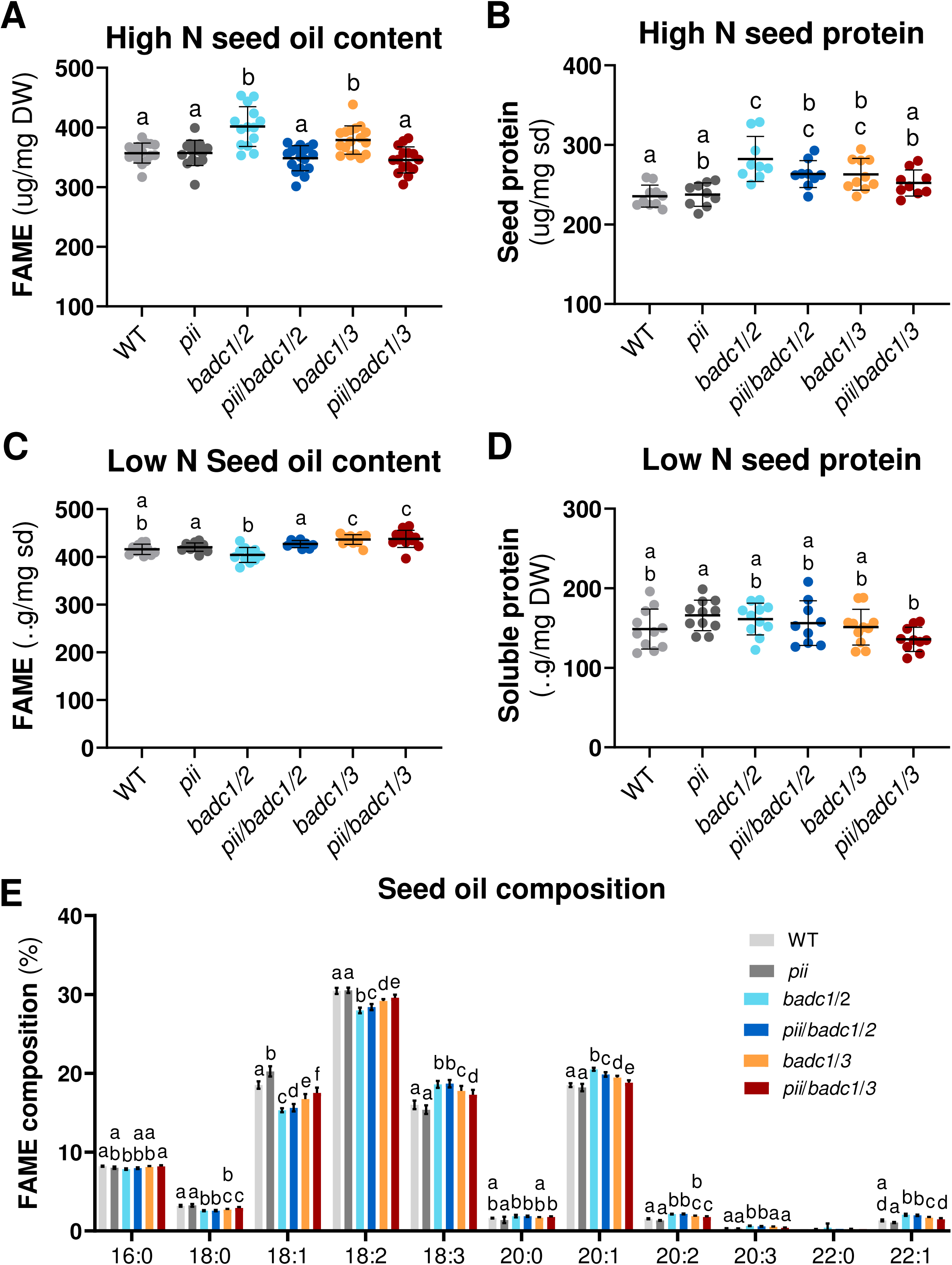
Comparison of seed oil and protein composition between *badc1/2 and badc1/3* double mutants their respective *pii/badc* triple mutants. Storage oil and protein were measured from plant lines fertilized with modified Hoaglands solution containing high N (100mg plant) (A, B) or no added N (C, D). Fatty acid composition of seeds was also compared from well fertilized seeds (E). Fatty acid composition of no added N seeds is found in Supplemental figure 4. Changes in seed oil and protein in *badc* and *pii* mutant combinations and wild type (WT) are presented as mean ± SD, statistical differences were analyzed by ANOVA and Holm-Šidák multiple comparison test. Different letters above each line represent significant differences in means (p-value < 0.05).

Many plant species demonstrate an inverse relationship between oil and protein in seeds, therefore we also analyzed protein content of seeds of each line. On the contrary, seed protein was increased in both *badc1/2* and *badc1/3* mutants by up to 18% (Fig. 5B) consistent with recent multi-omic analysis of just *badc1/3* mutant seeds (Kataya et al., 2024). Analogous to the seed oil results, the *pii/badc* triple mutant seed protein levels were also reduced when compared to their respective double mutants, with levels between those of the double mutant and WT (Fig. 5B). However, the seed oil and protein phenotypes were suppressed in *badc* double mutants grown under low N conditions (Fig. 5C, D). These results support that the *badc* double mutant seed oil and protein phenotypes are dependent on active PII (due to low 2-OG concentrations during high N) as N starved plants would be expected to have high levels of 2-OG and minimal PII function (Selim et al., 2020). When the fatty acid profiles of each mutant were examined, *badc1/2* and *badc1/3* demonstrated significantly higher levels of 18:3 and ≥20C fatty acids, at the expense of 18:1 and 18:2 in well fertilized plants (Fig. 5E). The corresponding *pii/badc1/badc3* triple mutant (Fig. 5E) partially complemented *badc1/3* parent line indicating the effects of the *badc1/3* double mutant on fatty acid composition may be PII dependent. The composition of *badc1/2* and its corresponding triple mutant showed similar trends, however, changes were quantitatively modest. In N starved lines the fatty acid composition was unchanged when compared to well fertilized plants (Fig. S4) which suggests that changes in seed fatty acid composition (rather than total oil amount) are N independent. Together, changes in seed oil and protein amounts support the conclusion that BADC proteins have regulatory control over both seed protein and oil accumulation, and both are dependent on the presence of PII.

### Acetyl-CoA carboxylase subunit stoichiometry is altered in *badc* double mutant lines

While there is clear evidence that loss of BADC proteins influences seed oil and protein accumulation, it is not clear what effect the loss of BADC and PII proteins may have on the stoichiometry of other htACCase catalytic and non-catalytic subunits. The AQUA-MRM technique that was previously utilized to quantify absolute amounts of htACCase subunits including BADCs in WT Arabidopsis seeds (Wilson and Thelen, 2018) was extended to include PII and was applied to developing seeds of each double and triple mutant background (Fig. 6). Surprisingly, in the WT and BADC double mutant backgrounds PII is more abundant than all BADC and BCCP isoforms combined (Fig. 6A). The *badc1/2* background (only BADC3 remaining) demonstrated an increase in levels of BADC3 and the α-CT subunit of the carboxyl transferase subunit and a slight increase in the levels of all other htACCase subunits except BCCP2 suggesting an increase in total HtACCase holoenzyme (Fig. 6B). Levels of PII were statistically unchanged but showed a mean increase greater than the increase of BADC3 (97 vs 56 fmol/µg protein, respectively, p=0.02; Fig. 6B). Thus, the increase in seed oil is linked not only to removal of BADC1/2 but also potentially to an increase in total active ACCase levels and corresponding PII. Conversely, in the *pii/badc1/2* triple mutant the levels of all remaining subunits were reverted back to WT, or below (Fig. 6B). These results mimic the effect on seed oil content in both *badc1/2* and *pii/badc1/2* (Fig. 5a), respectively, and may suggest that ACCase enzyme levels and stoichiometry contribute to the lipid synthesis and accumulation phenotypes of *badc1/2* and *pii/badc1/2*.

**Figure 6:**
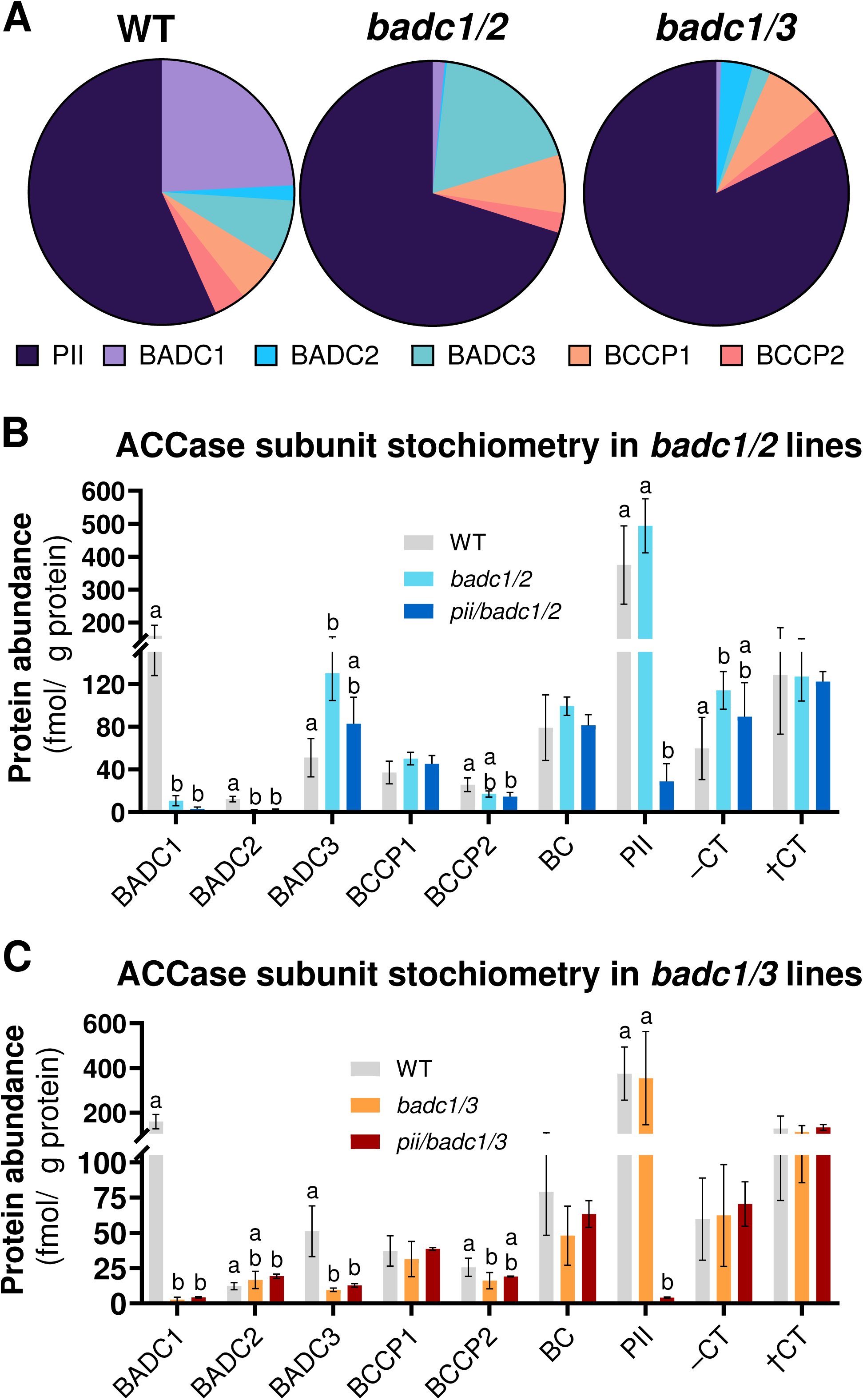
Determination of Acetyl-CoA carboxylase subunit abundance in *badc* double and *pii/badc* triple mutant developing seeds. Relative levels of PII as well as BADC and BCCP proteins are compared in 10-11 day after flowering developing seeds of wild type (WT) and *badc* double mutants (A). Changes in the absolute abundance and stoichiometric ratio of individual htACCase subunits were determined in *badc1/2* and *pii/badc1/2* (B); and *badc1/3* and *pii/badc1/3* (C) mutant combinations and compared to that of WT. Data is presented as mean ± SD, statistical differences were analyzed by ANOVA and Holm-Šidák multiple comparison test. Different letters above each line represent significant differences in means (p-value < 0.05).

In *badc1/3* double mutant lines no significant changes in the levels of htACCase subunits were seen with the exception of a decrease in BCCP2 (Fig. 6C). The results support that loss of BADC1 and BADC3 may have led to a modest negative feedback regulation of the biotin carboxylase sub-complex limiting total htACCase activity, or that post-translational control dominates regulation in *badc1/3* mutants (Fig. 6C). Further analysis of the *pii/badc1/3* triple mutant demonstrated that like *badc1/2*, levels of BCCP2 were partially complemented by the *pii* mutation (Fig. 6C). Seed oil was increased in *badc1/3* mutants (Fig. 5A) despite the limited effect on htACCase enzyme abundance (Fig. 6C) which suggests that the enhanced fatty acid synthesis in each of the *badc* double mutant lines are likely related to the effect of the remaining BADC2 or BADC3 on htACCase activity, the interaction of the remaining htACCase holo-enzyme with PII, post-translational control, and/or compensating mechanisms that affect partitioning of carbon into lipid biosynthesis. Levels of CTI proteins were also measured in developing seeds but were not changed (Fig. S5), suggesting that changes in seed oil content were not the result of the known α-CT regulation mechanism. Overall, the altered seed oil accumulation of the *badc* double and stacked *pii badc* triple mutants are not only due to the changes in htACCase activity through the loss of specific BADCs but also due to compensating regulatory changes that have affected total htACCase catalytic and regulatory subunit levels, of which appear to be distinct between the *badc1/2* and *badc1/3* backgrounds.

### PII demonstrates interactions with both BADC and BCCP proteins

Prior immunoprecipitation of htACCase utilizing a PII affinity column identified BCCP1 and BCCP2 as PII interactors, this was originally hypothesized to be a result of PII interaction with the biotin motif (Bourrellier et al., 2010). All three BADCs were also identified as potential PII binding proteins by immunoprecipitation, however, it was unclear if PII binds directly to BADCs or as a complex with BCCP proteins. To ascertain whether PII can interact independently with the individual subunits of ACCase, split-ubiquitin yeast two-hybrid protein-protein interaction assays were performed with PII as the bait and both catalytic and regulatory ACCase subunits (BC, BCCP1 or 2, BADC1, 2, 3) were utilized as prey. To ensure proper folding of these chloroplast proteins in the yeast cytosol, all proteins were expressed without chloroplast localization signals. Interaction was determined by serial dilutions (10-fold) of yeast expressing bait and prey grown on synthetic dropout media lacking tryptophan, leucine, histidine, and adenine (Fig. 7A), and by quantitative galactosidase reporter assays (Fig. 7B). Based on results from both analyses, a strong protein-protein interaction was found with the known PII binding partner NAGK (positive control), and clear PII interactions were demonstrated with BADC1, BCCP2, and BADC3. Minimal to no interaction with PII was indicated by BCCP1, BADC2, and BC. The selective media growth of yeast co-expressing PII and BCCP1/2 protein fusions (Fig. 7A) support previously published interaction of PII with BCCP proteins by immunoprecipitation (Feria Bourrellier et al., 2010a). Interestingly, these results suggest stronger interactions with BCCP2 than with BCCP1 (Fig. 7A, B). Within the BADC family, BADC1 showed the most robust binding/growth, followed by BADC3, whereas minimal to no interaction was evident with BADC2. Together these findings imply that PII may have preferential binding partners among the BADC and BCCP proteins, which may contribute to the differential phenotypes among the *pii/badc1/2* and *pii/badc1/3* lines.

**Figure 7:**
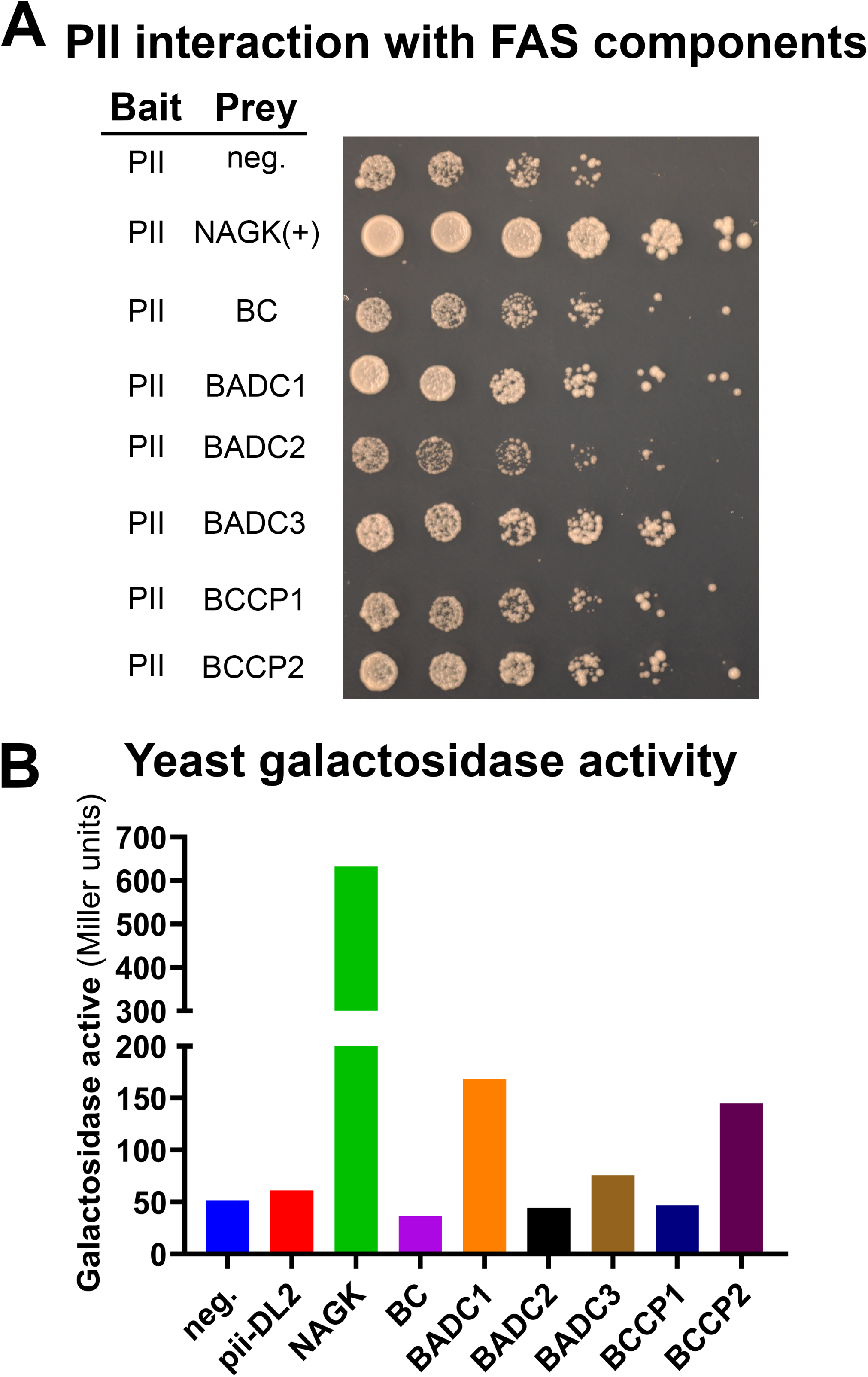
Yeast-two-Hybrid (Y2H) analysis of PII interaction with subunits of heteromeric Acetyl CoA carboxylase. (a) Serial dilutions of NYM51 yeast colonies expressing the pairs of indicated proteins grown on minimal media (SD -trp, leu, his, ade) for four days. From left to right each spot represents 10-fold serial dilution of a fresh culture OD_600_ 0.5. (b) Galactosidase assay measuring the activity of beta-Galactosidase in freshly cultured yeast expressing either control (CCW, DL2) or PII bait and indicated prey vectors. For both assays NAGK was used as a positive control for PII interaction while DL2 was used as a negative control.

## Discussion

In Arabidopsis and many dicots, fatty acid synthesis is a complex process that is mediated by a host of factors including substrate-product relationships, cellular conditions, and physical regulation through the activity of regulatory proteins. These proteins include the BADCs and PII which are believed to have no direct catalytic function but have been posited to alter the activity of the htACCase biotin carboxylase complex through their binding (Salie and Thelen, 2016). Correspondingly, seed oil accumulation and fatty acid composition in Arabidopsis is altered in both *badc* and *pii* mutants (Baud et al., 2010; Keereetaweep et al., 2018). While both PII and the BADCs were initially described as negative regulators, evidence in the literature and our own results suggest that regulation of the biotin carboxylase half-reaction of htACCase is more complex. Here we present evidence that both *badc1/2* and *badc1/3* mutants affect lipid and nitrogen metabolism, and that the effects on N and C metabolism are dependent on PII such that most of the *pii/badc* triple mutant phenotypes trend back toward wild type (Fig. 1-6). Interestingly, the relative phenotypes between each respective double and triple mutant combination are not always equivalent in each measure, demonstrating the differential phenotypes are also dependent on the remaining BADC isoform (BADC2 or BADC3) within each line. Additionally, through absolute quantitative proteomics we demonstrate that PII is more abundant than the BADC and BCCP isoforms combined, and each *badc* double mutant had a differential compensating effect on htACCase subunit levels in developing seeds, which again trended back toward wild type with the additional loss of PII (Fig. 6). Together, these results support the conclusion that the regulatory roles of each BADC isoform and PII on carbon and nitrogen metabolism are overlapping and distinct, which is further supported by the differential binding of BADC and BCCP subunits of htACCase to PII (Fig. 7). Figure 8 displays a proposed model of the combined regulation of htACCase by BADCs and PII, which is further discussed below.

**Figure 8:**
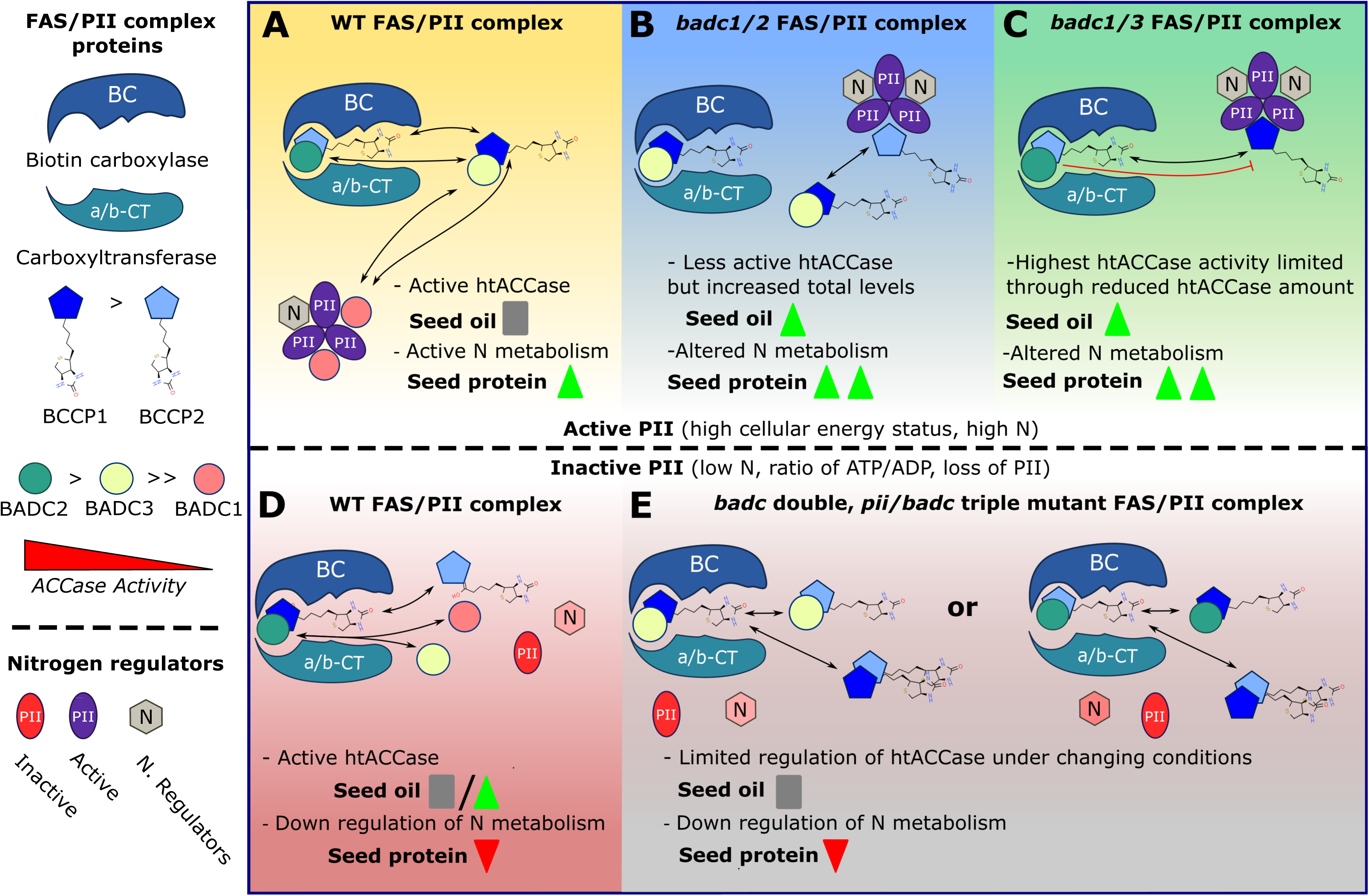
Model of BADC/PII regulation of htACCase and the interaction between N and C metabolism. The subunits of htACCase and their interaction with PII in wild type (WT), *badc* mutants, and *pii/badc* triple mutants. Resulting changes in Arabidopsis seed oil and protein are represented by green triangles (increase), red triangles (decrease), and grey bars (no change). Arrows represent possible substitutions of individual htACCase complexes.

### PII binds and independently regulates BADC and BCCP subunits of the ACCase complex to regulate fatty acid synthesis

The hypothesized inhibitor roles of the non-catalytic subunits of htACCase (BADCs and PII) were first described through *in vitro* assays where increasing amounts of BADCs or PII binding to ACCase reduced the *in vitro* activity (Bourrellier et al., 2010; Salie et al., 2016a). These roles appeared consistent with the mutant phenotypes of increased seed oil of the *badc* double mutants (Keereetaweep et al., 2018) and altered fatty acid composition of *pii*, suggesting unregulated fatty acid synthesis vs fatty acid elongation (Baud et al., 2010). Interestingly, the enhanced oil content of *badc1/3* over that of wild type was not present when grown under 24 hr light, suggesting that the enhancement of seed oil accumulation due to the *badc* double mutants is dependent on additional factors such as regulation of stromal energy status, pH, and cation concentration or changes in circadian rhythm (Ye et al., 2020b; Kim et al., 2023). Likewise, here we demonstrate that the oil phenotype of both *badc1/2* and *badc1/3* is also dependent on N availability (Fig. 5 A, C) further emphasizing the condition specific role of these regulators. The “negative regulator” hypothesis of BADC proteins became more nuanced with results that indicated BADCs may be required for organization of the quaternary structure of htACCase, such that holoenzymes that only contained BCCP but not BADC subunits had very low activity within *in vitro* buffer conditions optimized for ACCase activity (Shivaiah et al., 2020). The enhancement of ACCase activity was dependent on the BADC isoform with BADC2 followed by BADC3 showing the highest activity (Shivaiah et al., 2020). Interestingly, the addition of BADC1 improved ACCase activity modestly but this may be due to BADC1 sensitivity to pH, see Ye et al., (2020), or that physiologically BADC1 may function as a negative regulator. Thus, if BADCs are required for htACCase holoenzyme formation, then they are not simple activator/inhibitors but together the three individual BADCs may act as a biological rheostat allowing for varying levels of activity (Fig. 8).

Our results support and build upon this nuanced hypothesis by demonstrating that PII is able to independently bind to both BADC and BCCP proteins and most likely alters or inhibits their association with the BC subunit. First, protein-protein interaction studies in yeast demonstrate that apart from NAGK, PII can bind to both the essential BCCP and regulatory BADC proteins independent of the larger ACCase complex (Fig. 7), which is consistent with initial affinity chromatography which identified both BCCP and the then as of yet named BADC subunits (Feria Bourrellier et al., 2010a; Ye et al., 2020b). Additionally, PII demonstrated differential binding affinity for each BADC and BCCP isoform (Fig. 7) which suggests that PII may not function as a simple negative regulator of active htACCase. In developing WT seeds, PII is more abundant than all BADC and BCCP subunits combined (Fig. 6) indicating it could possibly sequester/exchange free BCCP and BADC isoforms based on its binding affinity and cellular energy and N status (PII binding is activated during a high energy and high N state (Selim et al., 2020)). PII may then function to differentially regulate which BCCP and BADC isoforms are within the htACCase holoenzyme in response to cellular energy and N status (Fig. 5, 7; (Smith et al., 2003; Feria Bourrellier et al., 2009; Feria Bourrellier et al., 2010a; Ye et al., 2020b)), through selective sequestration of BADC and BCCP proteins when active (Fig. 8A) or not modulating ACCase under low N and/or low cellular energy conditions (Fig. 8D).

We hypothesize that in both *badc* double mutant backgrounds the high oil phenotype reported here and previously (Keereetaweep et al., 2018; Ye et al., 2020b) is dependent on PII optimizing BADC-BCCP isoform ratios to fine tune ACCase activity under changing cellular energy, and nitrogen status (Fig. 5, 8A). The dependence on PII is supported by the *pii/badc1/2* and *pii/badc1/3* mutants that revert high oil phenotype of *badc* double mutant seeds to wild type (Fig. 5A, 8B, 8C), similar to low N (Fig. 5C, 8E) and constant light (Ye et al., 2020b). In a low N state 2-OG levels are high which inhibits the binding of PII to ACCase components (Bourrellier et al., 2010), analogous to the *pii* mutant (Fig. 8D). While plant growth under constant light limits the dynamic pH and ionic conditions associated with fluctuating energy status which are known to regulate both BADC and PII binding ability (Smith et al., 2003; Feria Bourrellier et al., 2009; Feria Bourrellier et al., 2010a; Ye et al., 2020b). Therefore, without PII modulation of the BCCP-BADC components of htACCase under changing conditions, the overall htACCase activity is reduced eliminating any enhancements of seed oil (Fig. 5). It should also be noted that *badc1/2* and *badc1/3* differ in their compensatory regulation of htACCase subunit stoichiometry (Fig. 6). Which along with the selective protein-protein interactions of PII with BADC-BCCP (Fig. 7), differential plant growth, protein, and lipid phenotypes (Fig. 1-5) supports that interaction with PII may be one of major differences between BADC isoforms. The selective interactions between PII and BADCs may lead to the differential phenotypes of the *badc1/2* and *badc1/3* mutants which have only BADC3 or BADC2 remaining, respectively.

### Differential phenotypes of the *badc* double mutants are resultant of the remaining BADC isoform and PII’s affinity to the remaining BADC/BCCP subunits

The *badc* double mutants have similar increases in seed oil and protein content, and comparable small increases in seedling ACCase activity, but had differential effects on vegetative biomass lipid and protein content (Figure 1, 2, 3,). The differences in vegetative tissues as compared to seed tissue is likely due to seed function as a storage tissue for excess fatty acid and protein synthesis, whereas vegetative tissue must adapt and maintain homeostasis to unregulated ACCase and altered N metabolism. The differential phenotypes on the leaves of the *badc1/2* and *badc1/3* mutants are explained both by the remaining BADC isoform in each double mutant, the selective binding of PII to the remaining BADC and BCCP isoforms, and the differential expression of htACCase subunits (Fig. 6) which together regulate ACCase activity (Fig. 8).

In the *badc1/3* mutant only BADC2 remains and the biotin carboxylase complex is limited to BC, a BCCP subunit, and BADC2 (Fig. 8C, E). This combination has been hypothesized to be one of the most active BC-BCCP-BADC combinations (Shivaiah et al., 2020; Table S1), and the role of PII in modulating BCCP1/2-BADC2 ratios is confined to interaction with BCCP2 within the ACCase complex (Fig. 7). Consequently, the vegetative tissue of *badc1/3* plants had modest effects on plant development as well as leaf lipid and protein composition (Fig. 1-3), demonstrating that BADC2 is able to functionally complement the *badc1/3* mutation and/or that BADC2 plays a major role in regulating the ACCase complex in vegetative tissues. In at least Arabidopsis, BADC2 expression is co-regulated with genes involved in regulation of circadian rhythm which would supports its role in maintaining cellular homeostasis (Kim et al., 2023). The further loss of *pii* in the *badc1/3* mutants reduced plant development and ACCase activity to levels lower than either parent line alone (Fig. 1, 2), underpinning PII’s role in BCCP/BADC regulation and ACCase activity even with limited interacting partners. The importance of these interactions was confirmed in seeds where the increased oil content of *badc1/3* mutants is dependent on PII (Fig. 5A). Consequently, recent transcriptomics and proteomics analysis of the *badc1/3* developing seeds, also suggested changes in both beta oxidation and fatty synthesis possibly in response to overproduction of fatty acids (Kataya et al., 2024) which would be consistent with a positive role of BADC2 at physiological concentrations. Protein quantification also measured a compensating effect in developing seeds of the *badc1/3* mutant through the reduction of BCCP2 (Fig. 6C), possibly to limit fatty acid over production, which was then alleviated by further loss of PII. While *In vivo* levels of ACCase were not measured in leaf or root tissue, it is likely that differential adaptation to the loss of *badc* expression also led to the differences during vegetative tissue growth.

In the *badc1/2* mutant only BADC3 remains (Fig. 8B, E). BADC3 demonstrated moderate binding to PII (Fig. 7) and would likely produce biotin carboxylase complexes (BC:BCCP1/2:BCCP1/2; BC:BCCP1/2:BADC3) which theoretically have intermediate ACCase activity and may not fulfill cell requirements for fatty acid synthesis (Shivaiah et al., 2020; Table S1). Interestingly, in developing seeds α-CT levels (the limiting subunit of htACCase; Wang et al., 2022) increased in the *badc1/2* mutant, along with total BADC3 and PII (Fig 6) which may be a compensatory mechanism for the reduced activity of the BADC3 containing htACCase complex. The corresponding *badc1/2* tissue demonstrated reduced plant development, and leaf lipid content. However, it is still unclear if the presence of BADC3, the loss of BADC2, PII interactions with remaining ACCase subunits or a combination of all these interactions within the ACCase complex are the defining factors. Changes in leaf protein as a result of PII interaction with NAGK or as of yet characterized N processes have also led to altered lipid content in *badc1/2* lines (Fig. 2C). However, the ratio 16:3/18:3 ratio in leaf MGDG and DGDG of *badc1/2* plants were largely not PII dependent (Figure 3A, C, D), which supports that the altered fatty acid production and enrichment of the chloroplast localized prokaryotic pathway of lipid biosynthesis corresponds to the loss of BADC1/2 while changes in total fatty acid synthesis and accumulation may result from altered PII regulation of N and C metabolism (Fig. 2, 3). Curiously, *badc1/2* seeds demonstrated increased oil (Fig. 5A) along with increased abundance of ACCase subunits (Fig. 6C) suggesting that seeds may have a compensatory mechanism for moderate activity of the htACCase complex containing only BADC3. Subsequently, the return of ACCase levels in *pii/badc1/2* mutants support that PII may have had negative effects on htACCase complex assembly in *badc1/2* double mutants. Together, these results suggest that the individual phenotypes of the *badc1/2* and *badc1/3* double mutants are related to the remaining BADC isoform, the ability of PII to modulate ACCase activity through interactions with the remaining BADC and BCCP subunits, and the plant adaptation to those specific changes in ACCase enzyme activity within different tissues (Fig. 8). More importantly, the selective interactions between PII and individual BADC and BCCP proteins may be the basis for post-translational regulation of the biotin carboxylase subunit in response to changes both N and cellular energy status.

### BADC proteins may co-regulate N metabolism through feedback regulation to PII

In Arabidopsis, there is clear evidence that PII has an important role in cellular N transport and metabolism within the chloroplast. PII is a major binding partner of NAGK and required for enzyme function, which is thought to be a regulatory step in the production of the amino acid arginine (Ferrario-Méry et al., 2006). There is also evidence of PII regulation in intercellular N transport (Ferrario-Méry et al., 2005; Ferrario-Méry et al., 2008), however the mechanism of action has yet to be confirmed (Selim et al., 2020). The loss of *pii* alone seems to have modest and largely non-significant effects on both N and seed protein accumulation in response to N availability (Baud et al., 2010). By contrast, mutants of *badc1/2* and *badc1/3* show significant differences in leaf and/or seed protein when compared to WT (Fig. 2B, 5B, 5D). The corresponding *pii/badc* triple mutants demonstrated a relative decrease in seed protein that was not significantly different than WT or their respective double mutant *badc* background. Our results indicate that BADC and possibly BCCP may regulate N metabolism through competitive binding and/or inhibition of PII (e.g. co-sequestration of BADCs or PII from further interacting with their binding partners of C or N metabolism). Contextually, the loss of BADC1 one the most abundant BADC isoforms (Fig. 6) with strong interactions with PII (Fig. 7) may lead to a relative increase in free active PII. At a minimum, excess PII would positively effect NAGK and promote arginine biosynthesis while negatively effecting hypothesized regulation of nitrite transport processes (Ferrario-Méry et al., 2008; Feria Bourrellier et al., 2009). Our results support that in the seeds of *badc* double mutants, possible changes in N assimilation and amino acid synthesis led to an increase in storage protein accumulation, which is consistent with recent results reporting increased seed oil in just *badc1/3* (Kataya et al., 2024). Interestingly, analysis of vegetative tissue, showed a variance between *badc1/2* and *badc1/3*, where *badc1/2* demonstrated both changes in leaf lipid and protein composition and increased nitrite toxicity suggesting that at least chloroplast N and C transport and metabolism were altered (Fig. 2B, 3, 4). These phenotypes were reverted to WT in the corresponding *pii/badc1/2* triple mutant supporting that PII interaction with ACCase components, or lack thereof in the case of *badc1/2*, is a regulatory check and balance between the accumulation of oil and protein. Overall, changes in leaf and seed oil and protein, and nitrite assimilation suggest that the regulation of N/C metabolism may be in part dependent on the interaction and competitive sequestration of PII and BADC proteins, which together are able to inform the larger regulatory network.

## Conclusions

### BCCP, BADC, and PII proteins function through competitive protein:protein interactions affecting both the stoichiometry of ACCase components/activity and N metabolism

Mounting evidence increasingly suggests that the biotin carboxylase sub-complex of htACCase is regulated through individual subunit expression, the regulated binding of the BADC and BCCP subunits in response to stromal conditions, and the interaction of the BADC and BCCP proteins to the N/C effector protein PII (Baud et al., 2010; Salie et al., 2016a; Ye et al., 2020a). Our results support the conclusion that the competitive binding of PII to each BADC and BCCP affects the stoichiometry of ACCase components and levels of free PII which can have downstream effects on ACCase activity and N metabolism. Consequently, the binding affinity between PII and individual BADCs may be the distinguishing characteristic between individual BADC proteins, allowing for various ACCase complexes with different activities. Finally, our results demonstrate that the non-catalytic subunits of htACCase (BADCs) are more than just negative regulators of ACCase activity, also impacting N metabolism within the cell via direct interaction with the PII effector protein. Future studies are needed to increase knowledge of ACCase regulation. It is especially urgent to determine the affinity between PII, BCCP, and BADC proteins and their stoichiometry within active htACCase complexes under different cellular energy and N status conditions. This knowledge is required for a more detailed understanding of how non-catalytic proteins contribute to the regulation of this essential metabolic enzyme.

## Materials and methods

### Identification of Arabidopsis triple mutants

Various genetic resources were used to create *Arabidopsis thaliana* lines containing different stacked mutations in the genes of interest. Wild type (*c.v.* Columbia (Col-0)) and T-DNA insertion mutants of *GLB1* (*pii;* PIIS2, (Ferrario-Méry et al., 2005), double mutants of *badc1* and *badc2 (badc1/2)*, *badc1* and *badc3* (*badc1/3* (Keereetaweep et al., 2018) were used as parent lines to produce the corresponding *pii/badc* triple mutants. Successful crosses of the tDNA triple mutant of *pii/badc1/badc2* and *pii/badc1/badc3* were screened by PCR (see, Table S2). Alternative *pii/badc1/3* triple mutants were also created by CRISPR genome editing. Multiple self-splicing ribozyme-flanked single guide RNAs (Gao and Zhao, 2014) targeting BADC1 and BADC3 were combined and cloned into the NotI and SacII sites of plasmid vector K63 (Shockey, 2020) to form plasmid J168 (Fig. S6). The AtUBQ10promoter:gRNA:CaMV35Sterminator cassette from J168 was cloned into the AscI and AvrII sites of the YAOpromoter:Cas9 DsRed-selectable binary plasmid pE717 (Li et al., 2013; Yan et al., 2015; Shockey, 2020) to produce plasmid E812, which was introduced into *A. tumefaciens* and used to transform pii T-DNA single mutant lines by floral dip (Clough and Bent, 1998). Progression of gene editing in candidate triple mutant plants was monitored by PCR amplification of the targeted DNA regions (for primers see Table S2) followed by restriction digestion of the resulting amplicons for detection of insertions or deletions that arose from error prone nonhomologous end-joining (NHEJ) DNA repair occurring after Cas9 target site cleavage. In this case, NHEJ may be expected to destroy either DraIII and/or PvuII sites in BADC1 and AvrII and/or PstI sites near the protospacer adjacent motifs (PAMs) of the guide RNA target sites. Promising lines were propagated and monitored for complete editing as well as segregation away from the DsRed marker associated with the Cas9 T-DNA. The PCR amplicons from successfully edited and segregated triple mutant plants were sequenced to confirm null mutations (Fig. S7). Finally, oil composition was compared to that of the tDNA triple mutant to confirm the phenotype (Fig. S1).

### Growth conditions

Seeds were surface sterilized by incubating in 70% ethanol for 5 min, then 0.1% SDS and 0.5% sodium hypochlorite for 10 min, followed by five rinses in sterile distilled water and stratification at 4°C for 48 hours. Sterile stratified seeds were sown directly onto soil and grown under a 16 h photoperiod with light intensity of 125-150 µmol m^-2^ s^-1^ in a walk-in growth room held at 21-23 °C. Plants were grown in 8 cm x 8 cm square pots at a density of 1 plant per pot for seed yield, oil, and protein and 4 plants per pot for analysis of leaf lipids, and biochemical experiments. For root length, ACCase activity, and nitrite growth studies, sterile stratified seeds were plated on media containing 1 % sucrose 0.5x strength MS salts (Murashige and Skoog, 1962) and 0.8 % phyto-agar. For nitrite experiments N deficient media was used and supplemented with 10 mM potassium nitrite (KNO_2_). All plates were sealed with Micropore tape (3M, St. Paul Mn, USA), then placed in a chamber set to conditions described above and allowed to grow for 10 days.

### Protein:protein interaction analysis

Yeast two-hybrid protein/protein interaction studies were performed using the DUAL membrane split-ubiquitin Y2H Hybrid kit 3 (Dual systems Biotech, Schlieren, Switzerland). The protein coding sequence of *PII* (*GLB1*) transcript, without its chloroplast localization signal, was amplified by PCR (Table S3) and cloned into the dual *Sfi*I sites in bait plasmid pBT3-C to create a C-terminal Cub-LexA-VP16 fusion. Similarly, multiple prey sequences including BC, BADC 1-3, BCCP 1-2, and NAGK were amplified without their chloroplast localization signal and cloned into prey plasmid pPR3-N to create NubG-HA epitope fusions (Table S3). Plasmid combinations were transformed into *Saccharomyces cerevisiae* strain NYM51 using the S.c. Easycomp transformation kit (Invitrogen, Walthum, MA, USA) and plated onto solid ‘SD-LT’ synthetic media (2% w/v glucose, 0.67 % w/v yeast N base without amino acids, 2% bacto-agar, and yeast amino acid dropout mix) lacking tryptophan and leucine (Invitrogen). Following successful transformation, yeast containing both bait and prey constructs were diluted to an optical density of 0.2 (600 nm). Colonies from each transformation were cultured in liquid SD-LT then diluted to an O.D.600nm of 0.2. Five µl of these suspensions and 1:10 serial dilutions thereof were plated onto SD media lacking tryptophan, leucine, histidine and adenine (Invitrogen), and allowed to grow on plates for 72 hours. The VP16 and LexA transcriptional activators present in the bait plasmid fusions regulate the LacZ reporter gene in strain NYM51, which allows for quantitative assessment of protein:protein interactions by *in vitro* galactosidase activity assays. Galactosidase activity was measured using the yeast beta-galactosidase assay kit (Fischer Scientific, Hampton, NH, USA) following manufacturer’s instructions.

### Vegetative analysis

Root length was measured from 10 day-old seedlings grown on solid media plates as described above. After 10 days plant roots were imaged, and root length was measured using ImageJ (Schneider et al., 2012) by tracing the developing roots using built in free hand line tool. Rosette mass was measured in 3 week-old rosettes by first cutting the plant at the base of the epicotyl, then drying the entire rosette at 65 °C overnight. Dried rosettes were weighed using a microbalance and analyzed for changes in rosette dry mass.

Leaf lipids were analyzed from 3 week-old wild type and mutants of both *pii* and *badc* proteins by quenching an entire rosette in 2 mL of 85 °C isopropanol for 15 min in an 8 mL glass tube with PTFE-lined cap. Lipids and total chlorophyll were extracted using a method modified from Bligh and Dyer (1959). In short, samples were ground in a polytron and 4 mL of a 4:1 chloroform: methanol (vol/vol) mix was added followed by centrifugation at 2500x *g* for 5 min. The supernatant was transferred to a new 13 mL glass tube and the solvent extraction above was repeated with 6 mL 2:1 chloroform: methanol and centrifuged again. The supernatants were combined and 2 mL of 0.8% KCl was added, shaken, and organic and aqueous phases separated by centrifugation. The total extraction was dried down under N_2_ and resuspended in toluene containing +0.005% BHT. Neutral and polar lipids were separated, and collected from leaf lipid extracts using an Agilent 1260 Infinity HPLC system and attached fraction collector following the separation described in (Kotapati and Bates, 2020) with modifications as indicated in (Kotapati and Bates, 2021; Wang et al., 2022).

### Leaf lipid and seed oil quantification

Total leaf and lipid fractions dissolved in toluene were analyzed by derivatizing lipids to Fatty acid methyl esters (FAMEs) and analyzing the resulting FAMEs were separated by gas chromatography and flame ionization detection (GC-FID) following a method modified from (Li et al., 2006). First, 20 µg of a Tripentadecanoin (15:0) internal standard dissolved in 50 µL toluene and 1 mL of 2.5% (v/v) sulfuric acid in methanol were added to 8 mL glass vials with PTFE lined caps. Samples were heated to 85 °C for 50 minutes. Following derivatization, resultant FAMEs were separated by the addition of 1 mL 0.8% (w/v) KCl and 0.5 mL hexane and centrifugation for 5 min at 2000x *g*. For seed oil analysis, dried seeds of individual plants were imaged using a DSLR, photographs counted using the Gridfree seed counting software (Hu and Zhang, 2021), and seeds weighed. Total seed lipids were then derivatized to FAMEs by adding 40 µg of 15:0 TAG internal standard dissolved in 500 µL toluene and 1 mL of 5% sulfuric acid in methanol in 8 mL glass vials with PTFE lined caps. FAMEs were then extracted by the phase separation as described above, however, 1 mL 0.8% KCl and 1.5 mL hexane were used. FAMEs were quantified against the 15:0 internal standard using an Agilent model 7890 GC-FID and a DB HEAVYWAX UI column (Agilent, Santa Clara, CA, USA; 30 m length, 0.25 mm inner diameter, 0.25 µm film thickness). The GC-FID was run in split mode at 1:10 for leaf and 1:40 seed samples; both GC injector and flame ionization detector were held at 255 °C with a helium flow of 1.05 mL/min. GC oven temperature started at 140 °C ramping at 20 °C/min to 200°C, then at 5 °C/min to 260 °C, followed by a 3 min hold.

### Leaf and seed protein quantification

Total soluble protein was analyzed from 10 mg fresh leaf material or 3 mg of dry seeds placed in a 2.0 mL tube which and ground in a bead mill (30 Hz for 3 min, with a 2.4 mm metal bead). Extraction buffer (1.8 mL; 100 mM Tris-HCl, pH 8, 2.5% (w/v) SDS, 10% (v/v) glycerol, 100 µM PMSF; 1X protease inhibitor cocktail (Complete Mini EDTA-free; Rosche Diagnostics, Mannheim, DE)) was added and samples thoroughly homogenized with the bead mill (15 Hz for 1 min). Samples were then heated at 80 °C for 15 min and seed material was sedimented by centrifugation at 15,000 x *g* using a benchtop centrifuge for 10 min, the resulting supernatant was transferred to a new 1.5 mL tube. Protein was quantified from a 1:2 dilution of the protein extract using the Pierce BCA protein assay kit for microplates (Thermo Scientific, Rockford, IL, USA) according to the manufacturer’s protocol and a microplate spectrophotometer (Spectramax 384 plus, Molecular devices San Jose, CA, USA; absorbance at 562nm).

### ACCase activity measurements

Leaf ACCase activity was measured in the developing Arabidopsis seedlings grown on plates (see above) at 10 days after germination following a protocol described in Salie *et al*. (2016b). In detail, pairs of 10-day old seedlings were harvested and quickly transferred to a pre-chilled 1.5 mL tube containing 150 µL cold extraction buffer containing 20 mM TES pH 7.5, 10 % (v/v) glycerol, 5 mM EDTA, 1 mM Dithiothreitol (DTT), 2 mM Benzamidine, 2 mM PMSF, and 1% Triton X-100. Samples were ground with a micro-pestle on ice and spun down in a 4 °C centrifuge at 10,000 x *g* for 1 minute. Next, 20 µL aliquots were then transferred to 1.5 mL tube containing 30 µL of either sample (100 mM Tricine pH 8.2, 100 mM KCl, 4 mM MgCl_2_, 1 mM ATP, 0.5 mM acetyl-CoA, 10 µM Haloxyfop, 1 µM NaH^14^CO_3_ [55.5 KBq]) or control (sample assay reagent with acetyl-CoA omitted) assay buffers. The samples were briefly mixed and incubated at 28 °C for 15 minutes. Enzymatic reactions were stopped with the addition of 20 µL of 2 M HCl and the resulting solution was pipetted into a scintillation vial containing a 1 cm^2^ square of filter paper. Samples were heated at 80 °C to remove excess [^14^C]CO_2_, 5 mL of Ecoscint added (National Diagnostics, Atlanta, GA, USA), and the resulting samples counted using a liquid scintillation counter (TRI-CARB 4910TR, Perkin Elmer, Walthum, MA, USA).

### Analysis of seedlings grown on nitrite

50 Arabidopsis seeds were plated and grown on solid media containing 10mM nitrite (described above) were grown for 3 weeks under constant light (20-25 µmol m^-2^ s^-1^) and constant temperature 20-23 °C. At the end of 3 weeks seedlings were imaged and the number of surviving seedlings that were green seedlings and had developed past the 2-cotyledon stage were counted.

### Absolute quantitation of ACCase proteins

Protein extraction and quantitation was performed as described in Wilson & Thelen, (2018). Prior to mass spectrometry analysis, 10 μg of each sample was aliquoted and normalized to the same concentration (10 µL volume) in urea buffer. AQUA™ peptides were used to monitor the abundance levels of ACCase catalytic subunits (α-CT, β-CT, BC, BCCP1, and BCCP2) and regulatory subunits (BADC1, BADC2, BADC3, CTI1, CTI2, CTI3, and PII). A 12.5 µL aliquot of AQUA™ peptide working solution was spiked into the biological samples before tryptic digestion, totalizing 1.25 picomoles of each AQUA™ peptide per sample. Reduction was performed at 30°C for 30 minutes with 10 mM DTT in 10 mM ammonium bicarbonate, followed by alkylation with 40 mM Iodoacetamide (IAA) in 10 mM ammonium bicarbonate for 1 hour at room temperature in the dark. Digestion was carried out by adding trypsin twice at an enzyme-to-protein ratio of 1:12 and incubating at 37°C for 14 hours followed by an additional 6 hours (final ratio of 1:6). After digestion, tryptic peptides were stored at -20°C for two hours, lyophilized via a centrifugal evaporator, and stored at -80°C until mass spectrometry analysis.

Samples were resuspended in 125 µL of 0.1% (v/v) formic acid in water Optima® LC/MS, resulting in a final AQUA™ peptide concentration of 10 femtomoles µL-1 and protein digest concentration of 80 ng µL-1. An injection volume of 5 µL for each biological replicate was used. Liquid chromatography and mass spectrometry analyses were conducted using an Easy nLC™ 1200 system (Thermo Fisher, San Jose, CA) coupled to a TSQ Altis™ Plus mass spectrometer (Thermo Fisher, San Jose, CA) over a 32-minute gradient at a flow rate of 300 nL/min. Transitions were optimized using the Skyline MRM method (64-bit, version 23.1.0.380) (MacLean et al., 2010; Bereman et al., 2012), and integrated into a final, optimized scheduled multiple reaction monitoring (MRM) method with a 3-minute retention time window. The summed intensity for eight transitions of light and heavy peptides was extracted using Skyline software. All AQUA™ peptides were injected within the linear detection range as previously detailed (Wilson & Thelen, 2018). The mass spectrometry proteomics data have been deposited to the ProteomeXchange Consortium via the PRIDE [1] partner repository with the dataset identifier PXD057351

### Statistical Analysis

All results are expressed as the mean ± stand deviation. Statistical significance was analyzed using GraphPad Prism Version 10.3.0 using one-way ANOVA followed by a Holm-Šidák multiple comparison test or Welches t-tests for pairwise comparisons.

### Reviewer access details for proteomics data

Reviewers can access the dataset (Project accession: PXD057351) by logging in to the PRIDE website using the following credentials: Username: and Password: 7T4ei0rJHNLt.

## Acknowledgements

The authors would like to thank Nora M. Zander for her assistance in the analysis of seed material. This work was supported by the National Science Foundation under grant Nos. 1829365 and 2242822; the United States Department of Agriculture National Institute of Food and Agriculture grant No. 2023-67013-39022, Hatch Project No. 1015621 and Multi-State Project No. 1013013; and the U. S. Department of Energy, Office of Science, Office of Biological and Environmental Research, under award No. DE-SC0023142. This work was supported in part by the U.S. Department of Agriculture, Agricultural Research Service. Mention of trade names or commercial products is solely for the purpose of providing specific information and does not imply recommendation or endorsement by USDA. USDA is an equal opportunity provider and employer.

## Author Contributions

P.D.B and J.J.T. conceived the project and procured the funding. M.G.G. and J.S. produced transgenic material. All experiments with the exception of protein quantification were performed by M.G.G. Protein quantification was performed by G. L. J. under the supervision of J.J.T. The manuscript was written by M.G.G. and P.D.B. with input from all authors.

## Abbreviations

ACCase: acetyl-CoA carboxylase
htACCase: heteromeric ACCase
BC: biotin carboxylase
BCCP: biotin carboxyl carrier protein
BADC: biotin attachment domain-containing protein
α-CT: α-carboxyltransferase
β-CT: β-carboxyltransferase
FA: fatty acid
FAS: fatty acid synthase/synthesis
TAG: triacylglycerol

## References

Baud S. Seeds as oil factories. Plant Reprod. 2018:31(3):213–235. doi:10.1007/s00497-018-0325-6

Baud S, Feria Bourrellier AB, Azzopardi M, Berger A, Dechorgnat J, Daniel-Vedele F, Lepiniec L, Miquel M, Rochat C, Hodges M, Ferrario-Méry S. PII is induced by WRINKLED1 and fine-tunes fatty acid composition in seeds of *Arabidopsis thaliana*. Plant J. 2010:64(2):291–303. doi: 10.1111/j.1365-313X.2010.04332.x

Baud S, Wuillème S, To A, Rochat C, Lepiniec LC. Role of WRINKLED1 in the transcriptional regulation of glycolytic and fatty acid biosynthetic genes in Arabidopsis. Plant J. 2009:60(6):933–947. doi:10.1111/j.1365-313x.2009.04011.x

Bereman MS, MacLean B, Tomazela DM, Liebler DC, MacCoss MJ. The development of selected reaction monitoring methods for targeted proteomics via empirical refinement. Proteomics. 2012:12(8):1134–1141. doi:10.1002/pmic.201200042

Bligh EG, Dyer WJ. A Rapid Method Of Total Lipid Extraction And Purification. Can J Biochem Phys. 1959:37(8):911–917. doi:10.1139/o59-099%M 13671378

Bourrellier ABF, Valot B, Guillot A, Ambard-Bretteville F, Vidal J, Hodges M. (2010). Chloroplast acetyl-CoA carboxylase activity is 2-oxoglutarate-regulated by interaction of PII with the biotin carboxyl carrier subunit. Proc Natl Acad Sci USA. 2010:107(1):502–507. doi:10.1073/pnas.0910097107

Clough SJ, Bent AF. Floral dip: a simplified method for Agrobacterium-mediated transformation of *Arabidopsis thaliana*. Plant J. 1998:16(6), 735–743. doi:10.1046/j.1365-313x.1998.00343.x

Cronan JE. The Classical, Yet Controversial, First Enzyme of Lipid Synthesis: Escherichia coli Acetyl-CoA Carboxylase. Micro Biol Mol Biol R., 2021:85(3):e00032–00021. doi:10.1128/MMBR.00032-21

Fan J, Yan C, Xu C. Phospholipid:diacylglycerol acyltransferase-mediated triacylglycerol biosynthesis is crucial for protection against fatty acid-induced cell death in growing tissues of Arabidopsis. Plant J. 2013:76(6):930–942. doi:10.1111/tpj.12343

Feria Bourrellier AB, Valot B, Guillot A, Ambard-Bretteville F, Vidal J, Hodges M. Chloroplast acetyl-CoA carboxylase activity is 2-oxoglutarate-regulated by interaction of PII with the biotin carboxyl carrier subunit. Proc Natl Acad Sci USA. 2010:107(1):502–507. doi:10.1073/pnas.0910097107

Feria Bourrellier AB, Ferrario-Méry S, Vidal J, Hodges M. Metabolite regulation of the interaction between *Arabidopsis thaliana* PII and N-acetyl-l-glutamate kinase. Biochem Biophys Res Commun. 2009:387(4):700–704. 10.1016/j.bbrc.2009.07.088

Ferrario-Méry S, Meyer C, Hodges M. Chloroplast nitrite uptake is enhanced in Arabidopsis PII mutants. FEBS Lett, 2008:582(7):1061–1066. doi:10.1016/j.febslet.2008.02.056

Ferrario-Méry S, Besin E, Pichon O, Meyer C, Hodges M. The regulatory PII protein controls arginine biosynthesis in *Arabidopsis*. FEBS Letters. 2006:580(8):2015–2020. doi:10.1016/j.febslet.2006.02.075

Ferrario-Méry S, Bouvet M, Leleu O, Savino G, Hodges M, Meyer C. Physiological characterisation of Arabidopsis mutants affected in the expression of the putative regulatory protein PII. Planta. 2005:223(1):28–39. doi:10.1007/s00425-005-0063-5

Fokina O, Chellamuthu V-R, Forchhammer K, Zeth K. (2010). Mechanism of 2-oxoglutarate signaling by the *Synechococcus elongatus* PII signal transduction protein. Proc Natl Acad Sci USA. 2010:107(46):19760-19765. doi:10.1073/pnas.1007653107

Fokina O, Herrmann C, Forchhammer K. Signal-transduction protein PII from *Synechococcus elongatus* PCC 7942 senses low adenylate energy charge *in vitro*. Biochem J. 2011:440(1):147–156. doi:10.1042/bj20110536

Gao Y, Zhao Y. Self-processing of ribozyme-flanked RNAs into guide RNAs in vitro and in vivo for CRISPR-mediated genome editing. J Integr Plant Biol. 2014:56(4):343–349. doi:10.1111/jipb.12152

Gerhardt ECM, Parize E, Gravina F, Pontes FLD, Santos ARS, Araújo GAT, . . . Huergo LF. The Protein-Protein Interaction Network Reveals a Novel Role of the Signal Transduction Protein PII in the Control of c-di-GMP Homeostasis in Azospirillum brasilense. mSystems. 2020:5(6). doi:10.1128/mSystems.00817-20

Hsieh M-H, Lam H-M, van de Loo FJ, Coruzzi G.A PII-like protein in *Arabidopsis*: Putative role in nitrogen sensing. Proc Natl Acad Sci USA. 1998:95(23):13965–13970. doi:doi:10.1073/pnas.95.23.13965

Hu Y, Zhang Z. GridFree: a python package of imageanalysis for interactive grain counting and measuring. Plant Physiol. 2021:186(4):2239–2252. doi:10.1093/plphys/kiab226

Huerlimann R, Heimann K. Comprehensive guide to acetyl-carboxylases in algae. Crit Rev Biotechnol. 2013:33(1):49–65. doi:10.3109/07388551.2012.668671

Jasinski S, Chardon F, Nesi N, Lécureuil A, Guerche P. Improving seed oil and protein content in *Brassicaceae*: some new genetic insights from *Arabidopsis thaliana*. OCL, 2018:25(6):D603. doi:10.1051/ocl/2018047

Keereetaweep J, Liu H, Zhai Z, Shanklin J. Biotin Attachment Domain-Containing Proteins Irreversibly Inhibit Acetyl CoA Carboxylase. Plant Physiol, 2018:177(1):208–215. doi:10.1104/pp.18.00216

Kim S-C, Edgeworth KN, Nusinow DA, Wang X. Circadian clock factors regulate the first condensation reaction of fatty acid synthesis in Arabidopsis. Cell Reports. 2023:42(12).

Kataya, Amr & Nascimento, Jose & Xu, Chunhui & Garneau, Matthew & Koley, Somnath & Kimberlin, Athen & Mooney, Brian & Allen, Douglas & Bates, Philip & Koo, Abraham & Xu, Dong & Thelen, Jay. (2024). Comparative omics reveals unanticipated metabolic rearrangements in a high-oil mutant of plastid acetyl-CoA carboxylase. [Manuscript submitted for publication] Preprint: bioRxiv doi: 10.1101/2023.08.31.555777.

Kotapati HK, Bates PD. Normal phase HPLC method for combined separation of both polar and neutral lipid classes with application to lipid metabolic flux. J Chromatogr B Analyt Technol Biomed Life Sci. 2020:1145:122099. doi:10.1016/j.jchromb.2020.122099

Kotapati HK, Bates PD. ^14^C-Tracing of Lipid Metabolism. In Plant Lipids: Methods and Protocols, D. Bartels and P. Dörmann, eds (New York, NY: Springer US), 2021:59–80.

Krapp A. Plant nitrogen assimilation and its regulation: a complex puzzle with missing pieces. Curr Opin Plant Biol. 2015:25:115–122. 10.1016/j.pbi.2015.05.010

Li JF, Norville JE, Aach J, McCormack M, Zhang D, Bush J, . . . Sheen J. Multiplex and homologous recombination-mediated genome editing in Arabidopsis and Nicotiana benthamiana using guide RNA and Cas9. In Nat Biotechnol. 2013:31:688–691

Li Y, Beisson F, Pollard M, Ohlrogge J. Oil content of Arabidopsis seeds: The influence of seed anatomy, light and plant-to-plant variation. Phytochemistry. 2006:67(9):904–915. doi:10.1016/j.phytochem.2006.02.015

Li-Beisson Y, Shorrosh B, Beisson F, Andersson MX, Arondel V, Bates PD, Baud S, Bird D, Debono A, Durrett TP, Franke RB, Graham IA, Katayama K, Kelly AA, Larson T, Markham JE, Miquel M, Molina I, Nishida I, Rowland O, Samuels L, Schmid KM, Wada H, Welti R, Xu C, Zallot R, Ohlrogge J. Acyl-lipid metabolism. (epub: Arabidopsis Book). 2013:11:e0161. doi:10.1199/tab.0161

Liu H, Zhai Z, Kuczynski K, Keereetaweep J, Schwender J, Shanklin J. WRINKLED1 Regulates BIOTIN ATTACHMENT DOMAIN-CONTAINING Proteins that Inhibit Fatty Acid Synthesis. Plant Physiol. 2019:181(1):55–62. doi:10.1104/pp.19.00587

MacLean B, Tomazela DM, Shulman N, Chambers M, Finney GL, Frewen B, Kern R, Tabb DL, Liebler DC, MacCoss MJ. Skyline: an open source document editor for creating and analyzing targeted proteomics experiments. Bioinformatics. 2010:26(7):966–968. doi:10.1093/bioinformatics/btq054

Murashige T, Skoog F. A revised medium for rapid growth and bio assays with tobacco tissue cultures. Physiol Plantarum. 1962:15(3):473–497.

Oke OL. Nitrite Toxicity to Plants. Nature. 1966:212(5061):528–528. doi:10.1038/212528a0

Raboanatahiry N, Li H, Yu L, Li M. Rapeseed (*Brassica napus*): Processing, Utilization, and Genetic Improvement. Agronomy. 2021:11(9):1776. doi:10.3390/agronomy11091776

Rolletschek H, Schwender J, König C, Chapman KD, Romsdahl T, Lorenz C, Braun H-P, Denolf P, Van Audenhove K, Munz E, Heinzel N, Ortleb S, Rutten T, McCorkle S, Borysyuk T, Guendel A, Shi H, Vander Auwermeulen M, Bourot S, Borisjuk L. (2020). Cellular Plasticity in Response to Suppression of Storage Proteins in the Brassica napus Embryo. Plant Cell. 2020:32(7):2383-2401. doi:10.1105/tpc.19.00879

Sagun JV, Yadav UP, Alonso AP. Progress in understanding and improving oil content and quality in seeds. Front Plant Sci. 2023:14. doi:10.3389/fpls.2023.1116894

Salie MJ, Zhang N, Lancikova V, Xu D, Thelen JJ. A Family of Negative Regulators Targets the Committed Step of de Novo Fatty Acid Biosynthesis. Plant Cell. 2016:28(9):2312–2325. doi:10.1105/tpc.16.00317

Salie MJ, Thelen JJ. Regulation and structure of the heteromeric acetyl-CoA carboxylase. BBA-MOL CELL BIOL L. 2016:1861(9, Part B), 1207–1213. 10.1016/j.bbalip.2016.04.004

Schneider CA, Rasband WS, Eliceiri KW. NIH Image to ImageJ: 25 years of image analysis. Nat Methods. 2012:9(7):671–675. doi:10.1038/nmeth.2089

Schwender J, Hay JO. Predictive Modeling of Biomass Component Tradeoffs in Brassica napus Developing Oilseeds Based on in Silico Manipulation of Storage Metabolism. Plant Physiol. 2012:160(3), 1218–1236. doi:10.1104/pp.112.203927

Schwender J, Ohlrogge JB, Shachar-Hill Y. A Flux Model of Glycolysis and the Oxidative Pentosephosphate Pathway in Developing Brassica napus Embryos. J Biol Chem. 2003:278(32):29442–29453. doi:10.1074/jbc.M303432200

Selim KA, Ermilova E, Forchhammer K. From cyanobacteria to Archaeplastida: new evolutionary insights into PII signalling in the plant kingdom. New Phytol. 2020:227(3):722–731. doi:10.1111/nph.16492

Shivaiah KK, Ding G, Upton B, Nikolau BJ. Non-Catalytic Subunits Facilitate Quaternary Organization of Plastidic Acetyl-CoA Carboxylase. Plant Physiol. 2020:182(2):756–775. doi:10.1104/pp.19.01246

Shockey J. Gene editing in plants: assessing the variables through a simplified case study. Plant Mol Biol. 2020:103(1-2):75–89. doi:10.1007/s11103-020-00976-2

Smith CS, Morrice NA, Moorhead GBG. Lack of evidence for phosphorylation of *Arabidopsis thaliana* PII: implications for plastid carbon and nitrogen signaling. Biochim Biophys Acta. 2004:1699(1-2):145–154. doi:10.1016/j.bbapap.2004.02.009

Smith CS, Weljie AM, Moorhead GBG. Molecular properties of the putative nitrogen sensor PII from *Arabidopsis thaliana*. Plant J. 2003:33(2):353–360. doi:10.1046/j.1365-313x.2003.01634.x

So KKY, Duncan RW. Breeding Canola (Brassica napus L.) for Protein in Feed and Food. Plants. 2021:10(10):2220. doi:10.3390/plants10102220

Song H, Taylor DC, Zhang M. Bioengineering of Soybean Oil and Its Impact on Agronomic Traits. Int J Mol Sci. 2023:24(3):2256.

Su L, Wan S, Zhou J, Shao QS, Xing B. Transcriptional regulation of plant seed development. Physiol Plantarum. 2021:173(4):2013–2025. 10.1111/ppl.13548

Truan D, Huergo LF, Chubatsu LS, Merrick M, Li X-D, Winkler FK. A New PII Protein Structure Identifies the 2-Oxoglutarate Binding Site. J Mol Biol. 2010:400(3):531–539. 10.1016/j.jmb.2010.05.036

Vigeolas H, Waldeck P, Zank T, Geigenberger P. Increasing seed oil content in oil-seed rape (Brassica napus L.) by over-expression of a yeast glycerol-3-phosphate dehydrogenase under the control of a seed-specific promoter. Plant Biotechnol J. 2007:5(3):431–441. doi:10.1111/j.1467-7652.2007.00252.x

Wang M, Garneau MG, Poudel AN, Lamm D, Koo AJ, Bates PD, Thelen JJ. Overexpression of pea α-carboxyltransferase in Arabidopsis and camelina increases fatty acid synthesis leading to improved seed oil content. Plant J. 2022:110(4):1035–1046. 10.1111/tpj.15721

Wang S, Liu S, Wang J, Yokosho K, Zhou B, Yu YC, Liu Z, Frommer WB, Ma JF, Chen LQ, Guan Y, Shou H, Tian Z. Simultaneous changes in seed size, oil content and protein content driven by selection of SWEET homologues during soybean domestication. Natl Sci Rev. 2020:7(11):1776–1786. doi:10.1093/nsr/nwaa110

Wilson RS, Thelen JJ In Vivo Quantitative Monitoring of Subunit Stoichiometry for Metabolic Complexes. J Proteome Res. 2018:17(5):1773–1783. doi:10.1021/acs.jproteome.7b00756

Yan L, Wei S, Wu Y, Hu R, Li H, Yang W, Xie Q. High-Efficiency Genome Editing in Arabidopsis Using YAO Promoter-Driven CRISPR/Cas9 System. Mol Plant. 2015:8(12), 1820–1823. doi:10.1016/j.molp.2015.10.004

Yang Y, Kong Q, Lim ARQ, Lu S, Zhao H, Guo L, Yuan L, Ma W. Transcriptional regulation of oil biosynthesis in seed plants: Current understanding, applications, and perspectives. Plant Commun. 2022:3(5):100328. doi:10.1016/j.xplc.2022.100328

Ye Y, Nikovics K, To A, Lepiniec L, Fedosejevs ET, Van Doren SR, Baud S, Thelen JJ. The BADC and BCCP subunits of chloroplast acetyl-CoA carboxylase sense the pH changes of the light–dark cycle. J Biol Chem. 2020:295(29):9901–9916. doi:10.1074/jbc.RA120.012877

Ye Y, Fulcher YG, Sliman DJ, Day MT, Schroeder MJ, Koppisetti RK, Bates PD, Thelen JJ, Van Doren SR. Docking of acetyl-CoA carboxylase to the plastid envelope membrane attenuates fatty acid production in plants. Nature Commun. 2020:11(1). doi:10.1038/s41467-020-20014-5

